# The evolution of competitive effectiveness and tolerance in populations of *Drosophila melanogaster* adapted to chronic larval crowding at varying combinations of egg number and food volume

**DOI:** 10.1101/2023.07.24.550285

**Authors:** Srikant Venkitachalam, Chinmay Temura, Ramesh Kokile, Amitabh Joshi

**Affiliations:** Evolutionary and Organismal Biology Unit, Jawaharlal Nehru Centre for Advanced Scientific Research Jakkur, Bengaluru 560064 India

**Keywords:** density-dependent selection, competitive ability, pre-adult survivorship, pre-adult development time, dry weight per fly, dry biomass.

## Abstract

The theory of density-dependent selection posits that genotypic fitness can vary depending on the population density. Several long-term selection experiments on outbred populations of *Drosophila* adapted to chronically high larval densities have shown that the most common evolutionary response of such rearing is an increase in larval competitive ability. Some authors have proposed that a better understanding of the evolution of competitive ability can be achieved by its partitioning into effectiveness and tolerance components. Effectiveness is the amount of competitive inhibition imposed by a competitor on another, whereas tolerance is the degree to which a competitor can withstand inhibition. In this study, we have explored the evolution of effectiveness and tolerance components of competitive ability using three sets of outbred populations of *D*. *melanogaster* adapted to chronic larval crowding at different respective combinations of egg number, food volume and container dimensions. Effectiveness and tolerance were found to be dependent on the particular selection regime, the starting food amount and the trait used as the outcome of competition. Eclosion, dry biomass and dry weight per fly distributions over time indicated that competitive ability can also express itself in a time-dependent manner. The results suggest that larval competition and the evolution of competitive ability in *Drosophila* are extremely nuanced.

## Introduction

The theory of density-dependent selection is based on the notion that different genotypes or phenotypes may have a selective advantage at low vs. high population density conditions, and forms an important bridge between the fields of population genetics and population ecology (reviewed in Mueller 1997, 2009; Joshi et al. 2001). Some of the early ideas about the fitness of a type varying with population density were put forward by Charles Elton (Elton 1927), followed by Theodosius Dobzhansky (Dobzhansky 1950), J. B. S. Haldane (Haldane 1956), and Robert MacArthur (1962). The most well-known treatment of this concept was laid out in the book ‘The Theory of Island Biogeography’ by MacArthur and Wilson (1967). The authors defined *r-* and *K-*selection as the evolutionary consequences of selection under low and high population densities, respectively (MacArthur and Wilson 1967). Many models, both verbal and mathematical, stemmed from this work, and have been reviewed extensively (Mueller 1997; Joshi et al. 2001). The salient prediction coming from several of these models, including the one proposed by MacArthur and Wilson (1967), was that under *K*-selection, evolution of higher equilibrium population size *K* (also called ‘carrying capacity’ by some authors – critically reviewed in Mallet 2012) would occur (MacArthur 1962; MacArthur and Wilson 1967; Gadgil and Bossert 1970; Anderson 1971; Roughgarden 1971; Pianka 1972). Furthermore, the evolution of higher *K* would be consequent on the increase in the efficiency of resource utilisation of *K*-selected individuals (MacArthur and Wilson 1967; Pianka 1970). Over the last four decades, this prediction regarding efficiency was tested and subsequently rejected through a series of rigorous long-term selection experiments using populations of fruit flies *Drosophila melanogaster* adapted to high-density conditions (Mueller 1990; Joshi and Mueller 1996). These studies failed to find the evolution of greater food-to-biomass conversion efficiency as a correlate of selection for adaptation to chronically high population density or larval crowding.

Several alternate models of density-dependent selection predicted the evolution of increased competitive ability as a consequence of adaptation to high-density (Gill 1972, 1974; Asmussen 1983; Mueller 1988*a*; reviewed in Joshi et al. 2001; Mallet 2012). Some models also conflated or subsumed competitive ability under *K*-selection (MacArthur 1962; Pianka 1972; Gilpin et al. 1976; reviewed in Bradshaw and Holzapfel 1989; Joshi et al. 2001). Competitive ability, defined here as the sum of traits relevant to an individual’s performance in competition (Bakker 1961), could be equated to fitness in conditions of resource limitation (Prasad and Joshi 2003). The prediction that increased competitive ability evolves under adaptation to high-density was observed in several *Drosophila* selection studies, each of which saw the evolution of increased larval competitive ability for limited food (Mueller 1988*b*; Mueller et al. 1991). Moreover, some studies of adaptation to crowding in *Drosophila* saw an increase in larval feeding rate (Joshi and Mueller 1988, 1996), which had been shown to be a strong positive correlate of larval competitive ability (Burnet et al. 1977), leading eventually to a view that adaptation to larval crowding in *Drosophila* would almost invariably involve the evolution of increased larval feeding rate (Prasad and Joshi 2003).

In the last fifteen years, this canonical view of adaptation to larval crowding via increased larval feeding rate was challenged by three selection studies on adaptation to larval crowding in *Drosophila* species, including *D*. *melanogaster*, with the high densities implemented by varying combinations of egg amount, food volume and container dimensions (Nagarajan et al. 2016; Sarangi et al. 2016). In these studies, crowding-adapted populations did evolve greater larval competitive ability, but through increased efficiency with respect to development time, and not increased larval feeding rate as seen in the previous studies (Nagarajan et al. 2016; Sarangi et al. 2016).

What became apparent from these studies was that populations adapted to high larval densities at varying egg number, food amount and container dimensions actually experienced different ecologies in the culture vials, which likely altered their evolutionary trajectories toward increased larval competitive ability (Sarangi 2013, 2018; Sarangi et al. 2016). Thus, efficiency may or may not evolve in populations adapted to high-density conditions depending on the manner in which the high density was imposed. Moreover, the details of the high-density rearing also influenced the evolution of population stability parameters – with increased constancy stability evolving in populations subjected to selection for larval crowding in low food amounts (Dey et al. 2012; Pandey and Joshi 2022*a*), but not in populations selected for crowding with relatively higher food amounts, with higher food columns (Mueller et al. 2000; Pandey and Joshi 2022*b*). While the exploration of traits correlated with different types of high-density conditions is still underway (Sarangi 2018; Venkitachalam et al. 2022), the prediction regarding the evolution of increased competitive ability under chronic crowding has been directly or indirectly met in every *Drosophila* selection study seeking to test it (reviewed in Venkitachalam et al. 2022).

Here, we draw on the findings of the studies suggesting that there is more to crowding than just density (eggs per unit volume of food) as well as an earlier argument for thinking about the consequences of density dependent selection in terms of the effectiveness and tolerance components of competitive ability rather than just efficiency (Joshi et al. 2001), in order to see whether viewing the outcomes of adaptation to crowding experienced in different ways in terms of effectiveness and tolerance might provide deeper insights into the ecology and evolution of competitive ability in *Drosophila*.

We define effectiveness as the average degree of potential population growth inhibition imposed by individuals of a focal group on those of a competitor group, as compared to the inhibition imposed by competitor group individuals on each other. Tolerance is the average degree of reduction of such inhibition by the focal group, inflicted by the competitor group, as compared to the inhibition observed in the focal group individuals when competing amongst themselves. These definitions are very similar to those used by Joshi et al. (2001). This approach of splitting competition into a proactive and a reactive component has been discussed for several decades, albeit with varying terms being used for the two components.

Early work focused on components of competition in plant species – called aggressiveness (similar to effectiveness) and sensitivity (inverse of tolerance) (Breese and Hill 1973), or effect and response (Peart 1989; Goldberg and Landa 1991), respectively. Peart (1989) found little or no relationship between colonising ability (of seeds) and inhibition ability (of adult plants), and equated the terms to competitive effect and response, respectively. Goldberg and Landa (1991) found competitive effect and response to be uncorrelated in several species of herbaceous plants.

In *Drosophila*, some of the earliest studies on this concept were carried out by Mather and Caligari (1983). The authors studied competitive pressure (or aggressiveness) and response (used in lieu of sensitivity) in intraspecific larval competition between two strains of *Drosophila*, across three experiments, studying these aspects for both pre-adult survivorship and body weight of eclosed adults. While no difference was found in the response component of either strain, aggressiveness was found to differ among strains in all three experiments. The authors concluded that aggressiveness and response appeared to be independent in terms of their variation, and could combine additively to result in the competitive ability of a genotype (Mather and Caligari 1983). A continuation of the above-mentioned study found that competition in trio-cultures was a reflection of the competitive ability of each strain in duo-cultures (Caligari and Mather 1984). Paul Eggleston (Eggleston 1985) speculated that the aggressiveness and response of genotypes “must depend upon factors such as larval feeding rate, critical weights, food conversion efficiency, the secretion and excretion of various substances into the medium and even the exploitation of behavioural characteristics”.

An early selection experiment was also carried out on a *D*. *melanogaster* population, for low aggression and for high response (response here was equated with sensitivity, or the inverse of tolerance) – both markers of lowered competitive ability(Hemmat and Eggleston 1988). Both trajectories in response to selection led to a decrease in the average competitive ability of the population. However, selection for low aggression led to a clearer result than for high response (Hemmat and Eggleston 1988). This study, thus, showed that the two components of competitive ability could evolve independently, and this was explored in greater detail in a later study on interspecific competition (Joshi and Thompson 1995).

Joshi and Thompson (1995) tracked changes in competitive effectiveness and resistance (i.e., tolerance) over 11 generations in 3 sets of populations of *D*. *melanogaster* vs. *D. simulans*, competing in 3 different environments, respectively. At the end of the experiment, almost all population sets saw an increase in the overall competitive ability of *D. simulans*, initially the weaker competitor. However, this increased competitive ability was likely achieved via alternate routes in each population – different combinations of effectiveness and resistance were seen to evolve in each population (Joshi and Thompson 1995).

To summarise, the *Drosophila* studies on effectiveness and tolerance highlighted two things – a) that effectiveness and tolerance can be affected independently by ecological factors in a given population, and need not evolve together, and b) that effectiveness and tolerance can be defined using readouts of competition other than just survivorship, such as body size of adults. However, the limitations of some of these studies have been their reliance on inbred strains (which may greatly limit inferential reach: Rose 1984), or on using populations relatively naïve to larval competition (e.g., Mather and Caligari 1983; Eggleston 1985).

These limitations can be overcome by selection experiments such as those that established that competitive ability can evolve in outbred populations adapted to chronic larval crowding, and that this evolution can happen through the correlated evolution of different underlying traits, depending on the exact combination of egg number, food amount and container dimensions used to implement the larval crowding. However, despite the advances made in the understanding of effectiveness and tolerance in competitive ability, and their predicted usefulness in studying density-dependent selection, populations adapted to larval crowding experienced in different ways have yet to be examined for effectiveness and tolerance. In this study, we undertake a systematic examination of effectiveness and tolerance with regard to four different parameters that can be used as surrogates for evaluating larval competitive ability in several population sets of *D. melanogaster*, each adapted to larval crowding experienced in different ways. The measures of effectiveness and tolerance employed are similar to those used by Joshi and Thompson 1995 (see Methods).

Thee four parameters of competition we use – pre-adult survivorship, pre-adult development time, total dry biomass, and dry weight per fly – are each representative of the potential growth of a population. Pre-adult survivorship is the most straightforward readout (used in Bakker 1961; Mueller 1988*b*; Nagarajan et al. 2016; Sarangi et al. 2016) – only survivors to adulthood get a chance to reproduce and have possible non-zero fitness. Pre-adult development time may influence reproductive success in many ways (discussed in González-Candelas et al. 1990). In nature, the earliest developers can reproduce faster than late developers, and may get the advantages of a shorter generation time (Cole 1954). Even in discrete-generation laboratory cultures such as ours, where the generation time is fixed (see Methods), early ecloseing individuals may have greater opportunities for mating (reviewed in Mital et al. 2022). Moreover, crowded *Drosophila* cultures tend to accumulate metabolic waste over time (Botella et al. 1985; Borash et al. 1998). In such scenarios, early eclosing flies, assuming they also pupate earlier, may partly escape the toxic build-up that could otherwise prove lethal or reduce the capacity to reproduce in some way (Borash et al. 1998; Mueller and Barter 2015). Dry weight per fly is another potential measure of fitness – smaller flies often have some general loss of fitness (reviewed Mital et al. 2021), besides some mating-related disadvantages. Smaller males may have lower ability to manipulate females for mating (Mital et al. 2021), and females from crowded cultures, which are smaller in size than those from uncrowded cultures, may have lower fecundity than the latter (Pandey et al. 2022). It is, however, unknown if any variation in dry weight among flies within a crowded culture (as seen in Sarangi 2018, and in fig 5 of the current study) also results in size-based differences in fecundity. It may be reasonable to assume that the smallest flies from a crowded culture may also have the lowest fecundity, simply because they cannot provision the same number of eggs as larger flies. Additionally, there may not be much difference in the egg size of flies from crowded cultures compared to those from uncrowded cultures (Venkitachalam et al. 2022). Ideally, we would prefer to directly measure fecundity and count the resulting offspring produced by surviving flies in our competition studies, as was done by Gale (1964). However, due to logistical reasons we used indirect proxies such as dry weight per fly instead. The total dry biomass of eclosed flies from a culture is a result of both the dry weight per fly as well as pre-adult survivorship. Unlike the dry weight of individual flies, biomass data can be analysed more readily if some populations don’t show any eclosion (see fig 3, 4, 5 for examples). Biomass was earlier used as a readout of competition in several experiments by Bakker (1961, 1969). Finally, each of the four outcomes of competition used in the current study have previously been shown to be affected by larval crowding (Sang 1949; Bakker 1961; Ohnishi 1976).

## Methods

### Populations used

We used three sets of four replicate outbred *D*. *melanogaster* populations each, subjected to chronic larval crowding experienced at different combinations of egg number and food volume per rearing vial, along with the four ancestral control populations routinely maintained at moderate larval density. The four control MB populations are maintained at approx. 70 eggs in 6 mL cornmeal food, in cylindrical glass vials (8-dram vials: 2.2–2.4 cm inner diameter and approx. 9.5 cm height) (Nagarajan et al. 2016; Sarangi et al. 2016). Each replicate (*i* = 1 to 4) of all three sets of crowding- adapted populations is derived from replicate *i* of the MB set. This allows the implementation of a block design in the analysis of variance (see ‘statistical analysis’ section below). For all four sets of populations, matched replicate populations were assayed at the same time.

The details of the crowding-adapted population sets are as follows:

**MCU:** selected for larval crowding at 600 eggs in 1.5 mL food in the same vial type as MB. At the time of assaying, they had undergone selection for at least 229 generations. Each replicate population was assayed at a different generation – 229, 230, 231, 232 for replicates 4, 1, 3, 2 respectively.

**CCU:** selected for larval crowding at 1200 eggs in 3 mL food in the same vial type as MB. These are maintained at the same total eggs/food density as the MCU, but with twice the egg number and food volume. At the time of assaying, they had undergone at least 108 generations of selection.

**LCU:** selected for larval crowding at 1200 eggs in 6 mL food in slightly narrower 6-dram vials (see Sarangi 2018). This maintenance regime was somewhat similar to that used in earlier studies for the CU populations (Mueller et al. 1993). At the time of assaying, they had undergone at least 107 generations of selection.

Additionally, we used a common competitor population with an orange eye (OE) mutation as a distinct visual marker in duo-cultures (see below). The OE population was maintained at low density, in a manner similar to the control MB populations.

Extensive descriptions of the ancestry and maintenance of these populations can be found in recent publications (Sarangi et al. 2016; Sarangi 2018; Venkitachalam et al. 2022). In brief, all sixteen populations are maintained on a 21-day generation cycle. Eggs are collected into vials at the respective selection density on day 1. Once the flies eclose, they are transferred to Plexiglas cages (25 × 20 × 15 cm^3^). This transfer takes place daily from around day 8 to day 20 in crowding-adapted populations, owing to a prolonged eclosion period in crowded cultures (see fig. 3 for an example). In MB as well as OE, transfer occurs only once, on day 11, by which time almost all eclosions are over. On day 18, the flies of all populations are provided food plates with a generous supplement of a paste containing live yeast mixed with water and a few drops of glacial acetic acid. On day 20, they are provided a food plate for egg laying for a duration of about 18 hours. On day 21, the next generation commences, starting with a new round of egg collection.

### Standardisation

Prior to starting the experimental assay, we reared each population in a common environment to minimise potentially confounding effects of non-genetic inheritance. This was done by rearing each population in the MB type, low density regime of approx. 70 eggs in 6 mL cornmeal medium. The eggs of each population were collected in vials and reared for 11 days, and thereafter the eclosing flies were transferred to a Plexiglas cage. These populations were provided a Petri plate with cornmeal medium and a generous smear of live yeast paste made with water and a few drops of glacial acetic acid. After around 60 hours, we removed the yeast-laden plates and added a cornmeal plate with vertical surfaces provided for egg laying. This plate was provided for one hour, and served to remove fertile eggs previously incubated inside the female flies. This ensured that most eggs laid for the experiment were more or less freshly fertilised, and no artificial head starts were being provided to larvae from previously incubated eggs hatching several hours earlier than others. The assay was started after this step.

### Competition experiment: start

After the initial egg laying plate was removed, a fresh plate was added for 5 hours to each population’s cage. This was a harder plate for easier egg removal, made with 2% agar along with yeast and sugar (Venkitachalam et al. 2022). After 5 hours, each plate was removed from its respective cage. The eggs on the plate were transferred to a surface of 1% agar. We kept two cages for the OE population, owing to a larger requirement for eggs. The eggs from both the cages were mixed on the 1% agar surface after plate removal.

### Egg collection

At this stage, eggs were exactly counted up to the required amount using a soft bristle brush and collected in bunches for transfer to experimental vials. These 8-dram vials (same as the ones used to rear MB) contained either 1, 1.5 or 2 mL of cornmeal food. They were further divided into two categories: mono-culture, which received 400 eggs of a focal population or OE; and duo-culture, which received 200 eggs of a focal population and 200 eggs of OE. Thus, each vial had a total of 400 eggs, in either 1, 1.5 or 2 mL of food. There were five replicate vials per culture type, per food level, per selection regime, per block. We additionally set up low density vials with 70 eggs in 6 mL cornmeal food in a similar fashion. In this case, duo-culture vials had 35 eggs focal population + 35 eggs OE. Each replicate set (MB*i*, MCU*i*, CCU*i*, LCU*i*, *i* = 1…4), along with OE, was assayed together per maintenance generation. A total of 228,600 eggs, spread over 720 vials, were thus used for the experiment. After egg collection, the vials were kept at 25 °C in constant light in 70-90% relative humidity. The position of the racks (containing vials of various selection regime and density combinations) were shuffled daily to randomise the rearing environment as much as possible.

### Collection of eclosing flies: Pre-adult survivorship and development time measurements

On the 8^th^ day from egg collection, observations were started in order to collect eclosing flies from the culture vials. In each observation check, all vials were visually scanned for any eclosed flies. These checks were done 12 hours apart on the night of the 8^th^ day, morning of the 9^th^ day, night of the 9^th^ day and morning of the 10^th^ day, respectively. Afterwards, the checks were carried out 24 hours apart (10^th^ day, 11^th^ day, 12^th^ day…and so on). This was done until all eclosion ceased.

In each check, eclosed flies from each vial were transferred to a corresponding empty transfer vial. These flies were then frozen by dousing the transfer vials in liquid nitrogen for approx. 2 minutes. After this, flies from each vial were taken out onto a stereomicroscope platform for separation and counting. Flies from mono-cultures were segregated and counted on the basis of sex (female or male). Those from duo-cultures were segregated and counted on the basis of eye colour (wild type or orange) and sex. Any flies that inadvertently escaped during the transfer, freezing or counting process were noted (if noted before escape, sex and eye colour were recorded too).

For each vial, the pre-adult survivorship of a population was obtained by summing up the counts of all eclosed flies per eye colour (male and female) and taking its proportion to the total eggs of the respective population added into the culture. The pre-adult development time was taken as the average time of eclosion per eye colour, per sex.

After counting, we transferred the flies to 1.5 mL centrifuge tubes kept in respectively labelled sachets, in order to freeze and store them for subsequent weight measurements.

### Biomass and dry weight collection

Flies eclosing from each vial were transferred into their respective packages and stored at -20 °C for weighing later (after conclusion of the entirety of survivorship and development time assays). In these transfers, there were exceptions made as to the flies that could not be weighed, although they were noted for survivorship and development time. These exceptions included:

a. Any flies found stuck in food. These were discarded from weighing due to the likely presence of an additional confounding mass of leftover food.
b. Any flies that moved deep inside the cotton plug during eclosion, and got crushed in the process.
c. Any flies that were otherwise crushed during counting.
d. Any inadvertently escaped flies.
e. Any human error regarding sachet identity during fly transfer. The packet receiving the wrong transfer was discarded (see supplementary material for a detailed list of such exceptions).

These exceptions meant that the biomass was usually a small underestimate of the actual biomass (see supplementary material).

As an additional step for increased rigour, in order to control for any inadvertent transfers or losses, we also approximately re-counted all the flies from each tube before weighing. Any tube that had flies outside of ±15% of the number expected from survivorship counts was discarded from biomass readings. For dry weights, this percentage was relaxed to -30% of the expected counts for the lower limits, as lower than expected counts could still provide enough flies for obtaining the dry weight per fly. This discarding process resulted in some treatments having less than 5 replicate vials (at least 1 replicate vial was present in every case).

Flies were transferred according to their development time into one of three different time interval bins. These bins were:

**T1:** all flies eclosing up to day 11 (approx. 270 hours development time) were binned into the T1 sachets.

**T2:** all flies eclosing between day 11 and day 15 (approx. 270-370 hours development time) were assigned as T2.

**T3:** all flies eclosing after day 15 were kept in the T3 marked sachets. For T3, since the number of flies eclosing was generally lower, all vials were pooled together per combination of block × food level × population × culture type × sex.

The first time-interval bin, T1, was assigned such that it captured the vial duration of MB maintenance (which are transferred to cages on day 11), and could be classified as containing the early eclosing flies. The end of the second time bin, T2, completed a week of eclosion from the vials, and was set as such for logistical convenience. The bins are visually represented in the eclosion profiles in fig. 3.

### Dry biomass and dry weight measurement

Dry biomass measurements were taken to capture the weight of all eclosing flies from a particular centrifuge tube (containing a combination of block × food level × population × culture type × sex). Dry weight measurements were taken for a fixed sample of these flies, to obtain more accurate per fly weights, as was done in previous experiments (Sarangi 2018). By taking a constant number of flies per measurement, machine error per fly was also standardised. Thus, for dry weight per fly, a total of 10 flies (max.) were taken per reading. In case 10 or less flies were present in the tube, all the flies were weighed for both biomass and dry weight (lower limit set at 5 flies per tube). The weighing process was as follows:

1. Remove tube from freezer. Keep in convection oven (tube lid open) and dry at 70 °C for 36-42 hours.
2. Weigh the tube along with the flies stored within.
3. Remove all the flies and weigh the empty tube.
4. Add 10 flies randomly sampled from the removed flies, into the tube, and weigh again. (The weight of all flies + tube) – weight of tube = resulting dry biomass.

((The weight of 10 flies + tube) – weight of tube) ÷ 10 = resulting dry weight per fly.

### Effectiveness and Tolerance: calculations and properties

The measurement of both effectiveness and tolerance was broadly inspired by the definitions used by Joshi and Thompson (1995). The values were calculated as follows:

Effectiveness of a focal (i.e., MB, MCU, CCU, or LCU) population =

(Competitive outcome of OE in duo-culture vs. focal population - Competitive outcome of OE in mono-culture) ÷ Competitive outcome of OE in mono-culture

Tolerance of a focal population =

(Competitive outcome of focal population in duo-culture vs. OE - Competitive outcome of focal population in mono-culture) ÷ Competitive outcome of focal population in mono-culture Competitive outcome was represented by either pre-adult survivorship, pre-adult development time, total vial dry biomass of eclosing adults (pooling time windows and sex), or dry weight per fly at eclosion (averaged across time windows and sex). With the exception of pre-adult development time, a greater negative value of effectiveness implied greater competitive ability of the focal population. A population with high effectiveness would suppress the common competitor’s mean competitive outcome more than the common competitor could suppress its own outcome in competition. For development time, a lower absolute value confers greater fitness on average (see introduction). Thus, positive values of effectiveness for development time signify greater competitive ability of the focal population.

For tolerance, the patterns are reversed. For all outcomes except development time, a positive tolerance value suggests greater competitive ability. High tolerance values mean that the 200 larvae of the focal population had a more beneficial outcome when competing against 200 OE larvae than vs. 200 additional focal population larvae. As with effectiveness, pre-adult development time tolerance is inversely related to the competitive ability of the focal population. Negative tolerance values imply greater focal population competitive ability for development time.

### Statistical Analyses

We performed type 3 (mixed effects model) factorial ANOVA on both effectiveness and tolerance for pre-adult survivorship, pre-adult development time, dry weight per fly and total dry biomass, respectively. Selection was treated as a fixed factor with four levels (MB, MCU, CCU, LCU). Starting food volume was also considered a fixed factor with three levels (1 mL, 1.5 mL, 2 mL – 1 mL excluded for development time and dry weight per fly, see below). Block was a random factor with four levels, representing common ancestry and handling during assays (replicates 1-4). In addition, we also carried out ANOVA on the raw data for each trait, before calculating effectiveness or tolerance.

For pre-adult survivorship, the raw data analysis included 2 additional fixed factors as compared to effectiveness or tolerance for survivorship – these were eye colour (2 levels - wild type (MB, MCU, CCU or LCU) and orange eye) and culture type (2 levels – mono-culture and duo-culture). The survivorship data were also arcsine-square-root transformed to verify the statistical differences observed in the untransformed data. For pre-adult development time, there was no development time data in some blocks for orange eye at 1 mL in duo-culture vs. MCU and CCU, due to them having 0% survivorship. Thus, the 1 mL treatment was excluded from the raw data analysis as well as effectiveness analysis. The ANOVA for the raw data for pre-adult development time also included sex as a factor (2 levels – female, male), in addition to selection, starting food volume, block, eye colour and culture type.

For total biomass, as in pre-adult survivorship, the factors included selection, starting food volume (1 mL, 1.5 mL and 2 mL), block, eye colour and culture type. An additional fixed factor, the time window of eclosion, was also included, with three levels (T1, T2, T3, see text above). Female and male biomass were pooled per replicate vial.

Dry weight per fly was not analysed in its entirety due to several points of data being missing, due to lack of eclosion, at 1- and 1.5-mL food, in OE, at different time windows, in different sexes. Dry weight effectiveness analysis was carried out for the 1.5 mL and 2 mL cultures only.

All analyses were conducted in STATISTICA for windows (StatSoft 1995). Tukey’s HSD was used for post-hoc pairwise comparisons, calculated manually in each case. All results were plotted in R version 4.1.3 (R Core Team 2022) using the ggplot2, forcats and tidyverse packages (Wickham 2016; Wickham et al. 2019).

## Results

### Pre-adult Survivorship

The tolerance and effectiveness of the MB, MCU, CCU and LCU populations for pre-adult survivorship are shown in figures 1a and 1b, respectively. The raw survivorship values for the data of crowded cultures containing 400 eggs in 1-, 1.5- and 2-mL food, from which the effectiveness and tolerance were computed, are given in supp figure 1a-c, respectively.

**Figure 1:**
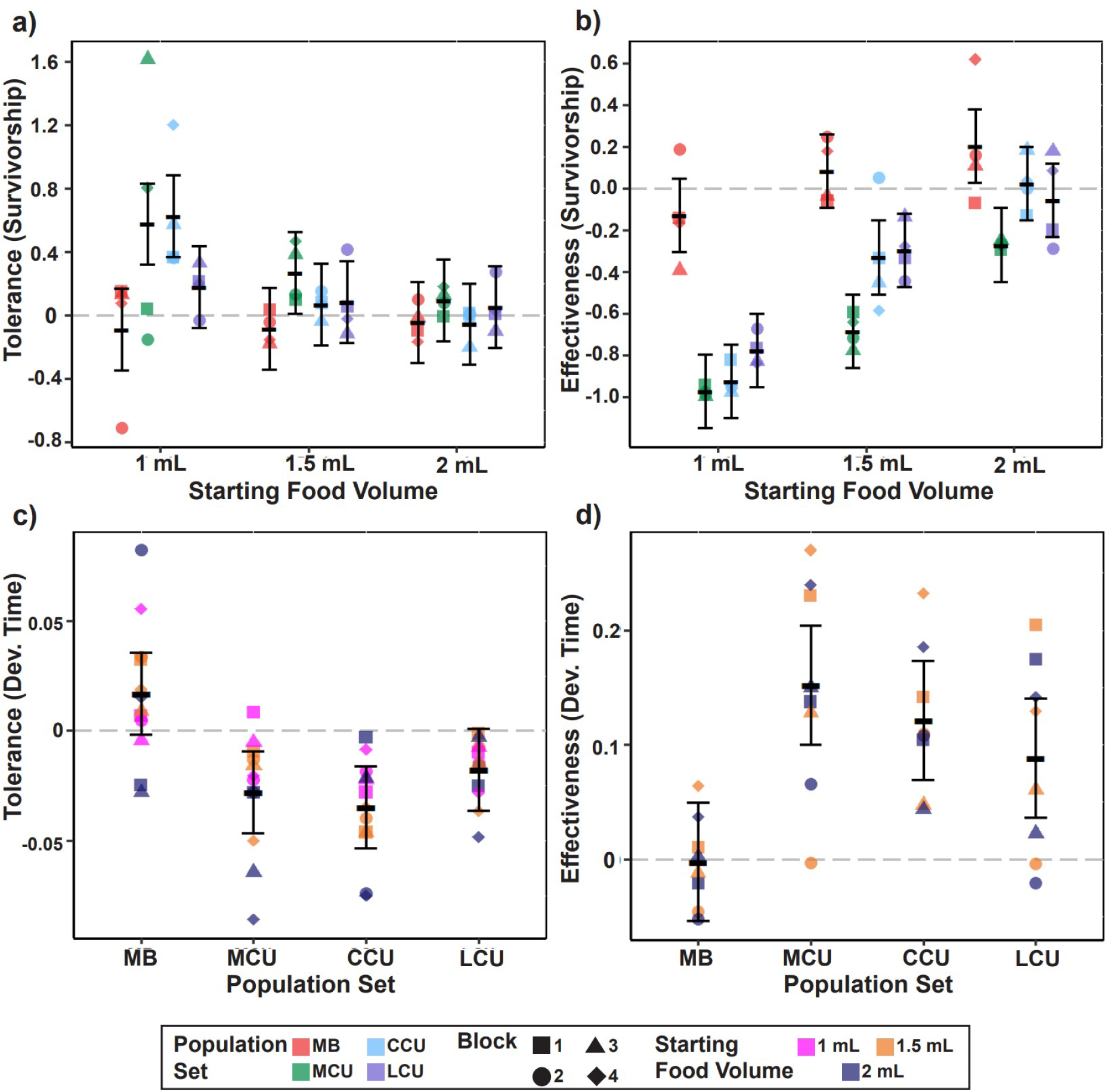
Tolerance and effectiveness of the three crowding-adapted population sets (MCU, LCU and CCU) as well as their low-density ancestral controls (MB). Error bars show ±95% confidence intervals around the mean of four replicate populations, based on the ANOVA. The individual plots are a) Tolerance for survivorship (interaction effect of starting food volume × selection, supp table 1); b) Effectiveness for survivorship (interaction effect of starting food volume × selection, supp table 2); c) tolerance for development time (main effect of selection, supp table 4); d) effectiveness for development time (main effect of selection, supp. table 5).

Increasing food level increased mean survivorship for all selection regimes and culture types (compared across supp fig 1a, 1b, 1c); main effect of food volume (*F*_2, 6_ = 33.80, *P* = 0.0005, supp table 3). The differences between selection regimes were the largest at 1 mL. Both MCU and CCU populations showed a gain in survivorship in duo-cultures as compared with mono-cultures. This was reflected in a mean tolerance significantly greater than 0 for both the population sets (fig 1a, interaction effect selection × food volume, *F*_6, 18_ = 3.15, *P* = 0.0273, supp table 1). CCU tolerance declined to values no different from 0 at both 1.5 mL and 2 mL food. However, MCU tolerance remained greater than zero when the starting food volume was increased to 1.5 mL food, although they too showed nearly zero mean tolerance at 2 mL food volume (fig 1a). Both MCU and CCU were significantly different from MB populations in mean tolerance at only 1 mL food. LCU tolerance values were, on average, between MB and CCU at 1 mL food, and nearly equal to CCU at higher food volumes (fig 1a). As a result, LCU tolerance was not different from MB, MCU or CCU at any food volume, nor was it significantly different from 0.

All three crowding-adapted population sets (MCU, CCU, LCU) had a significantly more severe competitive effect on the survivorship of the common competitor OE, than they did on themselves. This was reflected in the mean effectiveness data, with a significant main effect of selection (supp table 2, *F*_3, 9_ = 24.92, *P* = 0.0001). These patterns of differences were mainly driven by mean effectiveness at 1 and 1.5 mL food, with each of the MCU, CCU and LCU population sets differing significantly in effectiveness from the MB set, which was itself not significantly different from 0 effectiveness (supp table 2, interaction effect Selection × Food Volume, *F*_6, 18_ = 4.13, *P* = 0.0088). It is worth noting that the average effectiveness of both MCU and CCU at 1 mL food, likely the highest intensity of competition, was almost equal to -1 (fig 1b). This was due to OE survivorship against them being nearly 0% at the highest intensity of competition (supp fig. 1a). At 1.5 mL food, effectiveness for survivorship generally declined as compared to 1 mL, but this decline was more severe for CCU and LCU than for MCU, with both showing lower effectiveness than MCU at 1.5 mL food (fig 1b). MCU effectiveness was significantly negative even at 2 mL food – OE in duo-culture vs. MCU were performing poorly compared to OE in mono-culture at every single high-density food level. At 2 mL food, CCU and LCU effectiveness values were indistinguishable from 0. OE showed better survivorship in duo-culture vs. MB at 2 mL than in OE mono-culture, as can be seen from a positive effectiveness value for MB (fig 1b).

### Pre-adult development time (hours)

As mentioned in the methods section, the relationship of development time, all else being equal, tends to be inversely related to fitness – a greater development time implies lower fitness, on an average. Consequently, this also implies reversed patterns of tolerance (more negative value implies greater tolerance), as well as effectiveness (ascribed to greater positive values) for development time, as compared to survivorship.

Overall, we observed a pattern of increase of mean development time, as well as its variance, with food level, although these could not be tested for significance at the 1 mL food level due to lack of any surviving OE flies in duo-cultures vs. MCU and CCU.

Differences between mono- and duo-cultures tended to be larger at higher food levels for the crowding-adapted populations (fig3). However, the ANOVA did not show a significant interaction effect for selection × food volume for tolerance (supp table 4). There was a significant main effect of selection for tolerance (*F*_3, 9_ = 7.36, *P* = 0.0085, supp. table 4). Both MCU and CCU showed significantly negative tolerance (fig. 1c). They were also significantly different from MB, which was itself not significantly different from 0 tolerance. LCU showed no significant difference from any other selection regime, or from 0, for tolerance in development time (fig. 1c).

Effectiveness at 1 mL food could not be tested – the OE showed 0% pre-adult survivorship in competition against some MCU and CCU blocks, which meant that there was no development time to test in those cases. The ANOVA showed no significant main or interaction effect for starting food volume, implying an absence of statistical differences in effectiveness values at 1.5 and 2 mL food levels. The ANOVA did show a significant main effect of selection regime (*F*_3, 9_ = 8.05, *P* = 0.0064, supp. table 5), and the resulting pairwise comparisons are plotted in fig 1d, which pools both food levels per selection regime. All three sets of crowding-adapted populations were significantly more detrimental to OE than OE themselves, on average. This can be seen from the significantly ‘positive’ effectiveness values of MCU, CCU and LCU in fig 1d, whose means are not different from each other either. Average MB effectiveness was not significantly different from 0, nor was it significantly different from that of LCU, whereas both MCU and CCU had greater effectiveness than MB (fig. 1d).

Average variance for development time did not show any significant differences between mono- and duo-cultures (data not shown).

### Biomass

Average tolerance for biomass is plotted in fig. 2a. At the lowest food level, all three crowding adapted population sets (MCU, CCU, LCU) showed, on an average, increased biomass in competition with OE than in competition with themselves (interaction effect Selection × Food volume *F*_6, 18_ = 4.02, *P* = 0.0099, supp table 7). This was indicated by their significantly positive mean tolerance levels at 1 mL food (fig. 2a). All three population sets also showed higher tolerance than MB. CCU tolerance was significantly greater than LCU tolerance at 1 mL food (fig. 2a). At 1.5 mL food, both MCU and CCU maintained this pattern, but LCU tolerance dropped to an average that was positive, though not statistically different from 0, nor from MB tolerance. Finally, at 2 mL food, only MCU populations retained a positive mean tolerance, although this was not statistically different from MB (fig. 2a).

**Figure 2:**
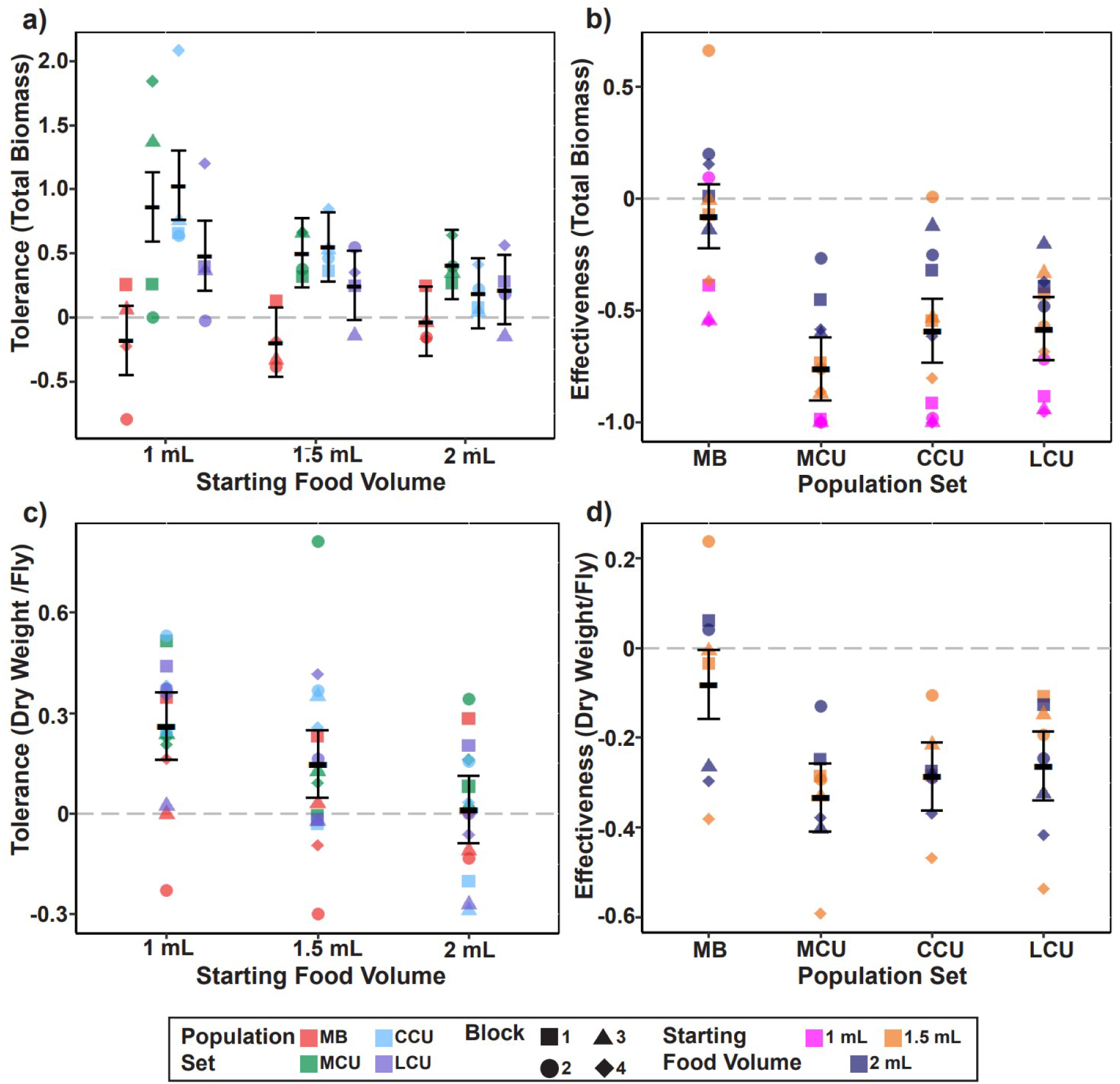
Tolerance and effectiveness of the three crowding adapted population sets (MCU, LCU and CCU) as well as their low-density ancestral controls (MB). Error bars show ±95% confidence intervals around the mean of four replicate populations, based on the ANOVA. The individual plots are a) Tolerance for dry biomass (interaction effect of starting food volume × selection, supp table 7); b) Effectiveness for dry biomass (main effect of selection, supp table 8); c) Tolerance for dry weight per fly (main effect of starting food volume, supp. table 10); d) Effectiveness for dry weight per fly (main effect of selection, supp. table 11).

For biomass-based effectiveness, only the main effects of selection and food level were significant in the ANOVA. The interaction between the two factors was not significant. All three crowding-adapted populations reduced OE biomass significantly compared to OE themselves, when averaged across all food levels (fig. 2b, *F*_3, 9_ = 20.98, *P* = 0.0002, supp. table 8). These were also different from MB, which was itself not significantly different from 0 in effectiveness. Additionally, all populations, on average, showed greater effectiveness at 1 mL food than at 1.5 and 2 mL food (*F*_2, 6_ = 41.32, *P* = 0.0003, supp table 8).

**Figure 3:**
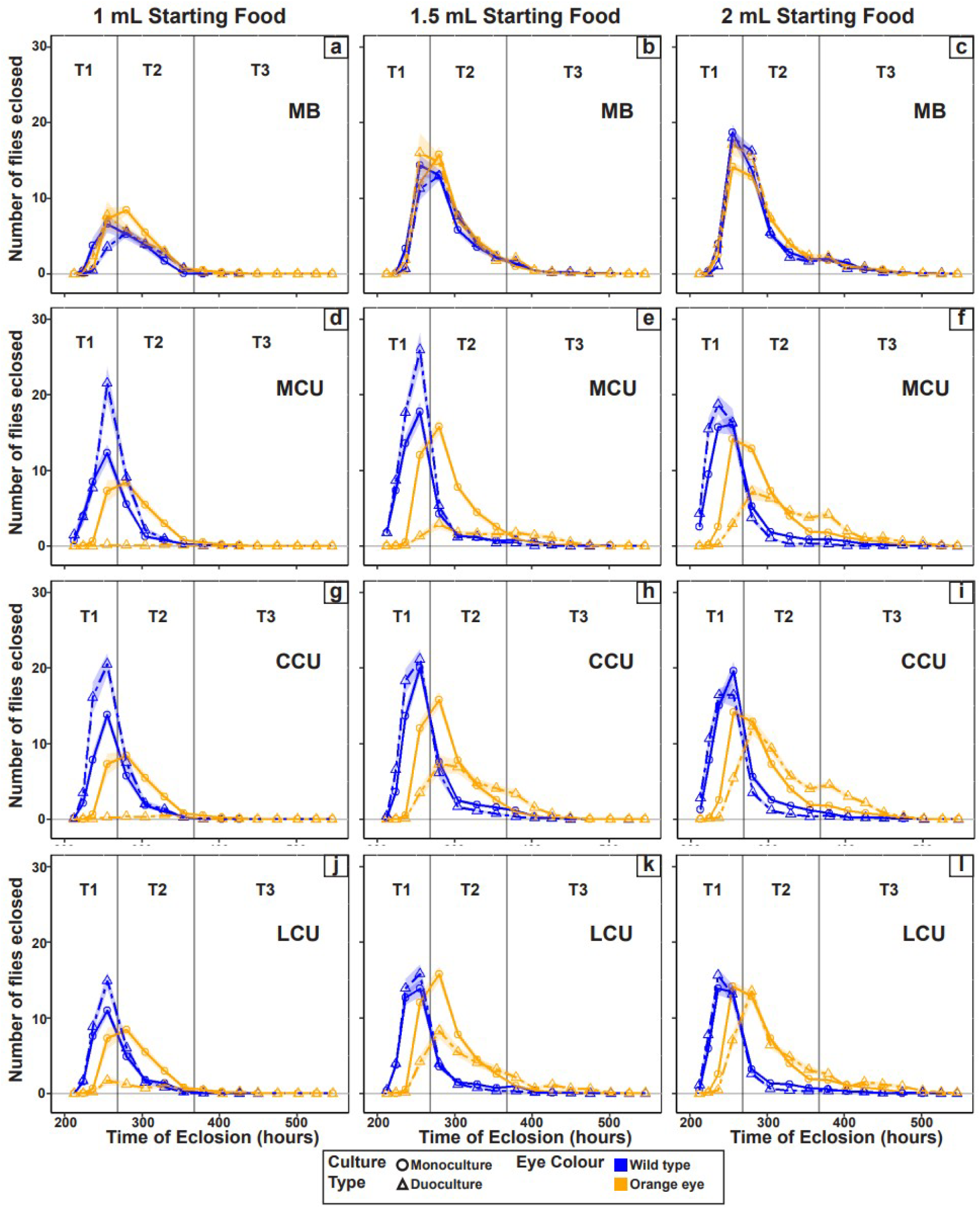
Eclosion over time for each focal population set across three starting food volumes (Top labels). Each row represents a population set; each column represents a starting food volume. Solid lines with circular points represent mono-culture (per 200 eggs); dashed lines with triangular points represent duo-culture. The shaded regions around the data points show S.E.M. The 3 time windows (T1, T2, T3; see methods) are also marked to aid visual interpretation of data in figures 4 and 5.

### Dry weight per fly at eclosion

The ANOVA for tolerance with respect to dry weight did not reveal any significant differences for selection regime (supp table 10). However, the patterns of gain in dry weight were largely similar to those seen for biomass (fig. 2a). Moreover, there was a significant main effect of starting food volume (*F*_2, 6_ = 7.11, *P* = 0.0262, table 10). Both 1 and 1.5 mL food had average tolerance greater than 0, which appeared to be driven mainly by the crowding-adapted populations (fig. 2c). There was nearly tolerance on average at 2 mL food, which was also significantly different from average tolerance at mL (fig. 2c).

Patterns for effectiveness of dry weight per fly were also similar to that seen in biomass. OE flies were smaller, on average, in competition against all three crowding adapted populations as compared to competition against OE (main effect of selection, *F*_3, 9_ = 8.76, *P* = 0.0049, supp. table 11). This can be seen from the non-zero effectiveness of MCU, CCU and LCU in fig 2d. Additionally, MB effectiveness was also significantly different from zero, implying that OE flies also lost dry body mass when competing against MB. This difference, however, was smaller than the reductions in dry body mass of OE flies when competing against the crowding-adapted populations – MCU, CCU as well as LCU were significantly more effective than MB in competition against OE (fig. 2d).

### Eclosion profiles

Figure 3 shows the profiles for eclosion of the different population sets across different culture types and starting food volumes. The number of eclosing flies is shown per 200 eggs in case of mono- cultures, in order to compare the values with the 200 eggs of the respective duo-cultures. Moreover, each time point on the X-axis is an average across replicate populations of the development time checks done over similar time windows per block.

MB and OE showed largely similar eclosion profiles over time (fig. 3a-c). The peak of eclosion increased in amplitude as food level increased, although this change was more pronounced from 1 mL to 1.5 mL than from 1.5 to 2 mL. Most 1 mL eclosions ended by the end of T2 time window, whereas they continued well into T3 at higher starting food volumes. There was a large gain in survivorship of OE at 2 mL food in duo-cultures when compared with mono-cultures (fig. 3c). This was also exemplified by the positive effectiveness for survivorship of MB at 2mL food (fig. 1b). MCU flies appeared to showed a relatively greater peak of eclosion in duo-cultures at both 1 mL and 1.5 mL food than in mono-cultures (fig. 3d-e). At 2 mL starting food (fig. 3f), the peak of MCU duo-culture eclosion was closer to mono-culture, and appeared lower than the peaks at 1 and 1.5 mL food. OE had zero (or very close to zero) survivorship against MCU at 1 and 1.5mL food duo-cultures, respectively (fig. 3d-e). At 2 mL food (fig. 3f), there were a steady number of OE duo-culture eclosions across a hundred hour period, although the relative peak of eclosion at mono-culture was greater. There was a significant difference between mono- and duo-cultures in the total eclosion of OE at 2 mL vs. MCU, however, as also reflected in negative effectiveness (fig. 1b). CCU eclosion profiles largely resembled MCU at 1mL (fig. 3d, 3g), as can also be seen from the tolerance and effectiveness plots (figures 1 and 2). At 1.5 mL food (fig. 3h), CCU populations lost the survivorship advantage in duo-culture vs. mono-culture (see fig. 1a), but continued to suppress OE survivorship (fig. 3h, 1b). Finally, at 2mL food (fig. 3i), survivorship differences between CCU mono- and duo-cultures disappeared, but a pattern for advantage in development time was present (fig. 1c). LCU populations followed a largely similar pattern to the CCU, with an overall smaller peak of eclosion, in both mono- and in duo- cultures (fig. 3j-l). There was also a small peak of eclosion that OE showed in duo-culture vs. LCU at 1mL food (fig. 3j) – a feature that was absent against the MCU and CCU populations.

### Dry biomass across time windows

The biomass measurements were divided across three time windows: T1, T2, T3 (up to 270 hours, 270-370 hours, post 370 hours from egg collection, respectively) (fig. 4). As in the eclosion profiles, mono-cultures are plotted per 200 eggs, to be matched with the 200 eggs seeded in the respective duo- cultures. These biomass values were also statistically tested for differences (supp table 9). There was a general pattern of biomass increasing with food volume (fig. 4). MB populations largely showed no differences from OE at any time point or food volume (fig. 4a-c). In duo-cultures, MCU populations consistently showed greater biomass than the mono-cultures at T1 across all food levels (fig. 4d-f) (fig. 4d). These large differences were also likely represented by a significantly positive total biomass tolerance of the MCU at each food volume (fig. 2a). At 2 mL, the T1 duo-culture MCU biomass was almost double that of the biomass of the (scaled to 200 eggs) mono-culture MCU populations (fig. 4f). Dry biomass in both T2 and T3 was significantly reduced compared to T1 across all food levels, in both mono- and duo-cultures in MCU (fig. 4d-f). The biomass of OE in duo-cultures vs. MCU was not significantly different from 0 at both 1 and 1.5 mL food (fig. 4d-e). Duo-culture biomass of OE vs. MCU was also significantly lower than the mono-culture OE at T1, at both 1.5 and 2 mL food (fig. 4e-f). However, in the most delayed time window from egg collection, T3, OE showed a non- significant pattern of greater biomass in duo-culture vs. MCU than in mono-cultures at 2 mL food (fig. 4f). In CCU, the patterns were largely similar to MCU (fig. 4g-i). There were some exceptions of note – the difference between duo-culture and mono-culture biomass of CCU at 1 mL, T1, appeared to be greater than the pattern seen in MCU(fig. 4g), and this was likely also reflected in the non- significant pattern of slightly higher tolerance for biomass in CCU vs. MCU at 1 mL food (fig. 2a).. However, the difference between the two culture types in CCU largely remained similar across food levels (fig. 4g-i), while this pattern of differences increased in the MCU (fig. 4d-f). Additionally, OE biomass in duo-culture was more comparable to mono-culture at both 1.5 mL as well as 2 mL food at T2 (fig. 4h-i). At T3 in 2 mL food, OE duo-culture biomass was nearly twice the value (scaled for 200 eggs) of the mono-culture biomass in competition against CCU, although this difference was not significant (fig. 4i).

**Figure 4:**
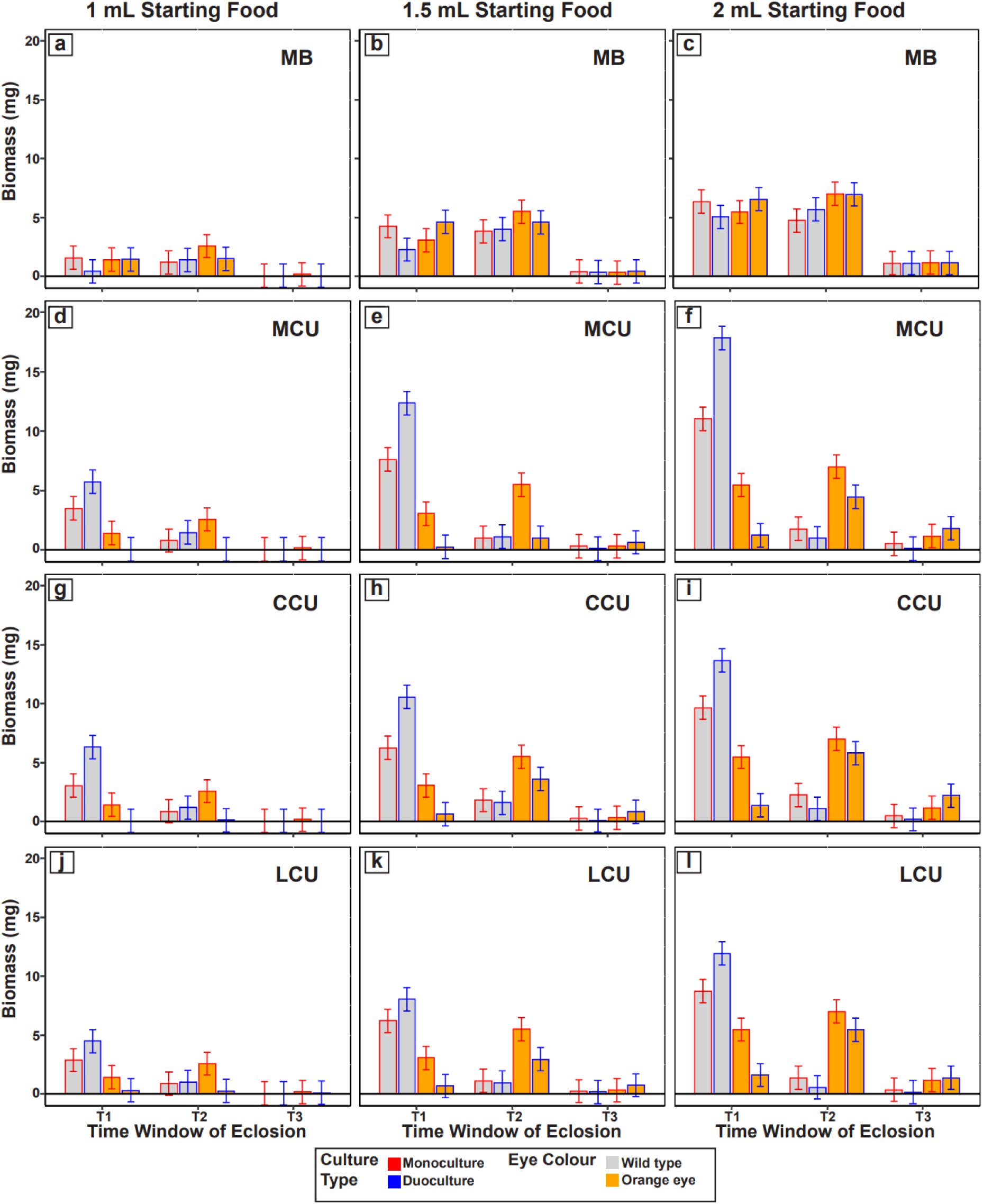
Biomass per population set per starting food volume, across the three time windows (T1, T2, T3; see methods). Error bars indicate ±95% Confidence Intervals around the means of four replicate populations (interaction effect selection × starting food volume × time window × culture type × eye colour, *F*_12, 36_ = 3.54, *P* = 0.0016, supp. table 9). Each figure label (a-l) corresponds to the same culture as labelled in figure 3.

LCU duo-culture biomass did not show much difference compared to that of mono-culture at both 1 mL and 1.5 mL food (fig. 4j-k). However, the duo-culture biomass at 2 mL was significantly greater than that of mono-culture at T1 (fig. 4l). The extent of this difference, however, was slightly lower than CCU, and much lower than MCU. T2 and T3 biomass of the LCU at 1.5 mL and 2 mL were not significantly different from 0 in duo-culture (fig. 4k-l). OE biomass values at duo-culture vs. LCU were (either significantly or with a non-significant pattern) reduced from those of mono-culture at both T1 and T2 across all food volumes (fig. 4j-l). It is also worth noting that there was very little difference in biomass of OE at T3 in duo-culture vs. LCU, as compared to the scaled biomass of OE in mono-culture (fig. 4l).

### Dry weight per fly across time windows

The dry weight per fly data could not be analysed in its entirety due to the lack of data for OE in 1 mL cultures across time steps (see fig. 1b, supp. fig. 1a). Nevertheless, the data are presented in figure 5 in order to see the overall patterns of dry weight over time, per population, per culture type. There were a few prominent patterns observable in the data. While OE in mono-cultures appeared larger than all focal population flies across all time windows, their dry weights in duo-cultures were reduced vs. the crowding adapted populations, especially in T1 (fig. 5d-l). In the dry weight distributions of 1 mL cultures, the flies eclosing in the T1 time window were generally larger than flies from T2 (fig. 5a, 5d, 5g, 5j). In the mono-culture distributions of 2 mL (fig. 5c, 5f, 5i, 5l), T1 flies as well as T3 eclosing flies appeared larger than T2 eclosing flies. This pattern changed however, in the duo-cultures of crowding-adapted populations. OE and (more prominently) MCU, CCU and LCU population flies were reduced or plateaued in size at T3 in duo-cultures as well as at T2, unlike in mono-cultures, which appeared to show an increase in average size from T2 to T3 (fig. 5c, 5f, 5i, 5l).

**Figure 5:**
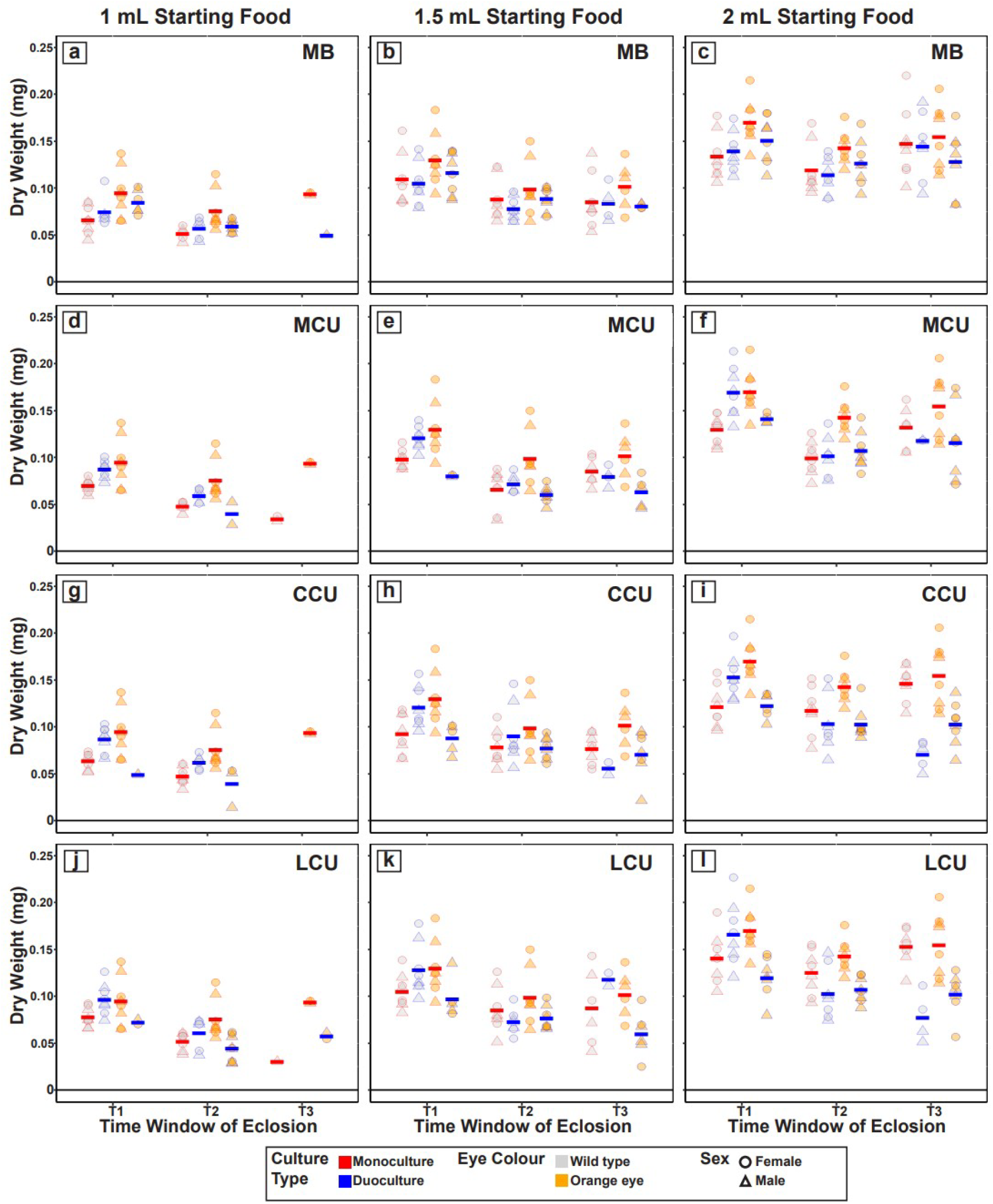
Dry weight per fly, for each combination of population set × starting food volume, across the three time windows (T1, T2, T3; see methods). Bold coloured bars represent mean under each category, respectively. Each figure label (a-l) corresponds to the same culture as labelled in figure 3.

### Low-density cultures

Data from the low-density cultures containing 70 eggs in 6 mL food are shown in supp fig. 3a-d. These show the potential values each of the traits used as outcomes of competition under crowding can take without the influence of crowding. Both mono- and duo-culture data are plotted.

## Discussion

Our results reinforce earlier findings that increased larval competitive ability is a consistent outcome of selection for adaptation to larval crowding. Moreover, our results highlight many nuances that become apparent upon splitting competitive ability along the axes of effectiveness and tolerance, for different measures of competitive outcome.

Starting food volume, even at relatively high densities, can be a major determinant of whether differences in competitive ability are seen between crowding-adapted and control populations. At 1 mL food volume, with a total density of 400 eggs/mL food, survivorship was lowest for all mono- cultures (supp fig 1). Furthermore, effectiveness and tolerance were most visibly different at this food volume for survivorship as well as biomass and dry weight per fly (fig. 1a-b and fig. 2). Out of the three crowding-adapted population sets, the MCU and CCU populations were practically identical in their effectiveness and tolerance at 1 mL along every axis of measurement (with perhaps CCU having a slight advantage in tolerance for development time (fig. 1c) as well as the biomass of early eclosing flies (figures 4d, 4g)). At higher food levels, however, CCU populations had relatively lower effectiveness and tolerance, and were similar to the LCU. In contrast, the MCU populations remained at average non-zero effectiveness and tolerance, at least at 1.5 mL food (figures 1 and 2). This is surprising, given that the MCU and CCU populations were similar in competitive ability at 1 mL food. However, the MCU populations had undergone over 100 more generations of selection than the CCU populations at the time of the current study. If we assume that the selection response has plateaued in both sets of populations, we should expect MCU and CCU to show similar competitive ability across all food levels. Alternatively, the MCU populations could, as they did against the LCU, show greater competitive ability than the CCU at every high-density condition examined. In contrast to both assumptions, the CCU populations competed just as well as the MCU when the total (eggs/food) density was close to their rearing density, but not when food levels were higher (and closer to the rearing food volumes of MCU). This suggests the involvement of some food column height effects in competitive ability as well.

LCU populations showed the lowest competitive ability in most conditions (as compared with MCU and CCU). This is not surprising, given that the LCU populations are reared at the lowest eggs/food density (200 eggs/mL for LCU vs. 400 eggs/mL in case of MCU and CCU). In the current experiment, the treatment containing 400 eggs in 2 mL food is the same total density as the one in which LCU is reared (200 eggs/mL food). Even at that treatment, however, LCU populations still performed, at best, on par with the MCU and CCU populations (figures 1 and 2). One possible exception was in the biomass of T3 (i.e., the last eclosing) OE flies, which showed a pattern of being the lowest against LCU at 2 mL duo-culture (fig. 4l). This might indicate the presence of greater effectiveness of the LCU larvae towards the later part of a culture, when any remaining food is likely riddled with high concentrations of metabolic waste (Borash et al. 1998). This is a scenario that is very likely to happen in the LCU native rearing cultures, wherein relatively high volumes of food are used. It remains to be seen if the LCU populations display any greater competitive advantages at the exact container dimensions, food volume and egg number quantities as their native rearing conditions (i.e., 1200 eggs, 6 mL food, 6 dram vials). While earlier experiments have examined competitive ability at the exact rearing conditions of LCU, they were too early in the selection process to give any clear results (Sarangi 2018). Moreover, it has recently been demonstrated by us that the density at the feeding band (a narrow volume of food, close to the surface, where the larvae feed) is an important determinant of the outcome of competition, more so than the total eggs/food density (S. Venkitachalam, V. S. Sajith and A. Joshi, ms in prep.). Given this knowledge, it cannot be ruled out that the LCU populations may have greater larval competitive ability close to the feeding band density at which they have been reared. Experiments are underway to determine if such specific competitive ability exists.

These results are also in agreement with the data of Pandey et al. (2022), who saw that MCU in mono- cultures had greater survivorship at higher densities of crowding compared to LCU. Moreover, the sensitivity of MCU survivorship to increased crowding was lower than LCU (Pandey et al. 2022). With the current study, we further add that MCU populations also show greater competitive ability across a wider range of crowding densities than LCU. In mono-cultures, Pandey et al. (2022) observed differences primarily at the highest density tested (300 eggs in 1 mL food). In contrast, in the current study, the effectiveness and tolerance of MCU show differences from LCU even at lower crowding densities. This indicates that larvae in duo-cultures can potentially show greater sensitivity to the outcomes of competition than mono-cultures.

Furthermore, while both average effectiveness and tolerance for pre-adult survivorship of each crowding-adapted population set changed similarly across crowding densities, there were some notable exceptions. MCU tolerance for survivorship was not significantly different from 0 at 2 mL food (fig. 1a), while the effectiveness was different (fig. 1b). This was true for CCU at 1.5 mL food (fig. 1a-b). LCU populations never showed a differencefrom 0, or from MB, in tolerance, but showed a difference in effectiveness (fig. 1a-b). The MB populations showed poor effectiveness at 2 mL but no change in tolerance (fig. 1a-b). These results together indicate that tolerance and effectiveness are likely independent in their evolution, as suggested previously (Mather and Caligari 1983; Caligari and Mather 1984; Eggleston 1985; Hemmat and Eggleston 1988; Joshi and Thompson 1995).

### The role of development time

While differences in survivorship and dry weight (and hence biomass) can be modelled to result from the simplest possible forms of exploitation competition (Bakker 1961; S. Venkitachalam and A. Joshi, unpublished), those in development time are likely to rise from more complex processes. In a crowded culture, there are two ways in which development time of a larva can be delayed – a) high concentrations of metabolic waste products building up and slowing developmental rate (Botella et al. 1985), and b) lack of space to feed, leading to lower overall feeding rate (discussed in Sang 1949; Bakker 1961). The phenomenon by which the development of a larva is arrested in a crowded culture, wherein it presumably faces severe competition and high concentrations of waste, has been termed ‘larval stop’ (Ménsua and Moya 1983). This process would be expected to lead to an extended development time distribution, as larval stop duration can be quite variable (Ménsua and Moya 1983). While the build-up of waste products such as urea, ammonia would be expected in all the food levels tested in the current study, the higher volumes of food would allow larvae to ingest more waste before running out of food, slowing overall development time, and possibly giving rise to larval stop (Botella et al. 1985). This could lead to a long tail of the development time distribution, which could further extend if more food were present, as larvae that were facing high levels of waste due to low initial feeding rate could still feed slowly in waste ridden food (see Borash et al. 1998). This is in overall agreement with what was observed in our study – while the 1 mL cultures showed a truncation in development time at around the T2 stage (fig. 3a, 3d, 3g, 3j), likely due to severe food shortage, the 2 mL cultures could sustain feeding, and thus eclosion, for a much longer duration – well into the T3 stage (fig. 3c, 3f, 3i, 3l). The crowded cultures with higher initial volumes of food also appeared to have a number of larvae feeding after the bulk of pupation was over, and had some moist food left over after all the eclosions were done (S. Venkitachalam, personal observation). This was in contrast to the 1 mL culture, in which food got depleted much faster and the remnants turned to dried powder by around day 10 from egg collection (S. Venkitachalam, personal observation). It is worth speculating here that the overall moisture of food in a culture can be an important factor in determining the volume of accessible food remaining in a culture. An additional noteworthy point is that the development times of every crowded culture are several hours longer than their respective low-density cultures (supp. fig. 2 vs. supp. fig. 3b), possibly suggesting some action of larval stop in each crowded food level. A previous study from our research group found the existence of some duration of larval stop in the MB populations, but not in the MCU populations (Pandey 2022).

The second possibility for delayed larval development has found some discussion in earlier papers – the existence of crowding for limited space, leading to reduced access to food, when a large number of larvae feed together in limited spatial constraints of the feeding band (Bakker 1961). A recent study by us also demonstrated that the larval density in the feeding band is a more important determinant of the outcome of competition than overall density (S. Venkitachalam and A. Joshi, in prep.). However, while some initial overcrowding of larvae may cause space shortages in the feeding band in the cultures used in the current study, these shortages are unlikely to be sustained to nearly the levels that are seen in the regular maintenance cultures of LCU or CCU, or even the MCU, which are each reared with higher absolute numbers of eggs. Thus, it is likely that the development time delays at moderately high feeding band densities, as used in the current study, are primarily through the consequences of metabolic waste build-up. This argument has some support from earlier studies as well, assayed at high feeding band densities (Borash et al. 1998; Mueller and Barter 2015). Given the argument developed above, it is also worth noting the pattern that 1 mL cultures appear to result in lower tolerance than 2 mL cultures (fig. 1c).

These results can also be compared with another study we conducted (S. Venkitachalam, A. Deep, S. Das and A. Joshi, ms in prep.), wherein MB, MCU, CCU and LCU populations were each competitively handicapped by providing a temporal head start in age to the OE eggs in competition against them (200 eggs focal population + 200 eggs OE in 2 mL food). When competing against each of the crowding-adapted population sets, head starts to OE gave greater development time advantages to OE adults against both CCU and LCU, than against MCU (S. Venkitachalam, A. Deep, S. Das and A. Joshi, ms in prep)… This was another example of MCU competitive superiority in 2 mL food volume, similar to the effectiveness and tolerance measurements in the current study, which once again suggests that the food column height at which the larvae of the population are adapted may play a major role in determining larval competitive ability in food columns of varying height.

### Distribution of dry weight over development time

The elongated development time distribution in crowded cultures with higher volumes of food also has consequences on the distribution of dry weight of eclosed flies. It has been shown before that in a crowded culture, early eclosing flies are usually larger than later eclosing flies (Hughes 1980; Sarangi 2018). This can be seen in the dry weight per fly distributions of 1 mL cultures, with T1 flies generally being larger than T2 flies (figure 5a, d, g, j). Sarangi (2018) also showed that in crowded cultures with higher food levels, early eclosing flies were larger than flies that eclosed in the middle of the distribution, but the flies eclosing towards the tail-end of the distribution were again quite large.

This was observed in the mono-culture distributions of 2 mL (fig. 5c, 5f, 5i, 5l), wherein T1 as well as T3 eclosing flies appeared larger than T2 eclosing flies. This pattern changed, however, in the duo- cultures of crowding-adapted populations (fig. 5f, i, l). Instead of showing an increase in size as in the mono-cultures, OE and (more prominently) focal population flies appeared to reduce (or plateau) in size at T3, compared to T2. A future experiment might explore this aspect in greater detail, with greater replication for vials and a greater resolution of binning in time windows.

Surprisingly, at 2 mL food, the smallest flies appeared to come from the crowding-adapted populations in duo-culture, in T3 (fig. 5f, 5i, 5l). Given the eclosion and biomass distributions (fig. 3 and 4, f, i, l, respectively), these were likely to be lower in number. However, this indicates the possible existence of crowding-induced late-eclosing variants in the MCU, CCU and LCU populations that can survive at smaller body masses than MB (see also Nagarajan et al. 2016, Sarangi et al. 2016).

### Fitness-functions – a matter of competitive ability distributions?

In the current study, the MCU, CCU and LCU populations had generally higher competitive ability than OE as seen by their increased effectiveness and tolerance across different starting food volumes (figures 1, 2) and the nuanced time-based expression of competitive ability displayed in biomass distributions (fig. 4). The duo-cultures of crowding-adapted populations also differed (or appeared to differ) from mono-cultures in eclosion, biomass and dry weight per fly distributions (figures 3, 4, 5). Given these two observations, we can speculate that the overall distribution and the variance of competitive ability likely determine the development time and dry weight distribution in crowded cultures. Duo-cultures containing eggs from crowding-adapted populations vs. OE are likely to furnish bimodal distributions in competitive ability, as opposed to unimodal distributions in mono- cultures, and this uniquely shapes dry weight, development time and biomass distributions in the crowded cultures.

This difference in competitive ability distributions becomes relevant when we compare interspecific and intraspecific competition experiments. In interspecific competition experiments (e.g., Moore 1952; Ayala 1969; Joshi and Thompson 1995) there have historically been large starting differences in competitive ability between the two competing species, indicating the presence of a bimodal starting distribution for competitive ability. This is in contrast to a selection experiment on intraspecific competition, which is more likely to start with a unimodal competitive ability distribution. Given the differences in competitive ability distributions, interspecific competition may result in very different patterns of survivorship, body size and development time for each species in the culture compared to their density-matched mono-culture counterparts. This would suggest that each competitor species could experience different underlying fitness-functions depending on whether they are in inter- or intraspecific competition. Consequently, these differences may lead to the evolution of very different traits for a given species in inter- vs. intraspecific competition, depending on the competitive ability of the species as well as the details of crowding under which the selection is carried out. In interspecific competition, different competitive strategies may be selected by the different species if there is more starting food, as there is likely to be enough survivorship of both species with a development time and size difference (as seen with MCU vs. OE in fig. 3-5). An interesting question also arises as to the rate of evolution of competitive ability (in the broad sense, as described in Joshi and Thompson 1996) in duo-species vs. mono-species systems.

## Conclusions

The evolution of competitive ability is a fundamental outcome of density-dependent selection in crowded conditions, and constitutes an important conceptual bridge between ecology and evolution. Our results indicate that outbred *D. melanogaster* populations can evolve increased competitive ability to different degrees depending on the precise nature of crowded rearing conditions. Furthermore, competitive ability, when split along the axes of effectiveness and tolerance, and measured for four different outcomes of competition, shows a staggering depth of nuance in its evolution, even in a seemingly simple laboratory culture system. Larvae of crowding-adapted populations can be superior competitors depending on the starting food volume, and show patterns of time-dependence in their competitive superiority, as seen from eclosion, biomass and dry weight per fly distributions over time (fig 3, 4, 5). While there have been some recent advances, both theoretical and experimental, in the study of density-dependent selection (Fronhofer et al. 2022; Bertram and Masel 2019; see also Than et al. 2020), our results highlight the need for understanding the ultimate consequences, as well as the mechanistic bases, of the depth of nuance visible in the process of competition and evolution of competitive ability. This requires greater exploration both in simple laboratory systems as used in the current study, as well as complex natural ones as discussed by other authors (Travis et al. 2013; Morimoto and Pietras 2020). Where logistically possible, the experiment design used in the current study can be further expanded in order to better study the ecology and evolution of larval competition. Such expansions include the use of multiple marked competitors (e.g., de Miranda et al. 1991; Santos et al. 1992), as well as more complex experiments incorporating replacement series or substitution methods (De Wit 1960; Seaton and Antonovics 1967; Mather and Caligari 1981).

## Acknowledgements

We thank Sajith V. S., Medha Rao, Rajanna N. and Muniraju for help with the experiments. S. Venkitachalam thanks Viveka Singh for help in logistical planning and re-checking of data. S. Venkitachalam and C. Temura were each supported by a doctoral fellowship from the Jawaharlal Nehru Centre for Advanced Scientific Research. This work was supported by a J. C. Bose National Fellowship from the Science and Engineering Research Board, Government of India, to A. Joshi, a Department of Biotechnology, Government of India - JNCASR project, “Life Science Research, Education & Training”, and, in part, by A. Joshi’s personal funds.

## Appendix: Supplementary Material

**Supp. Fig. 1.**
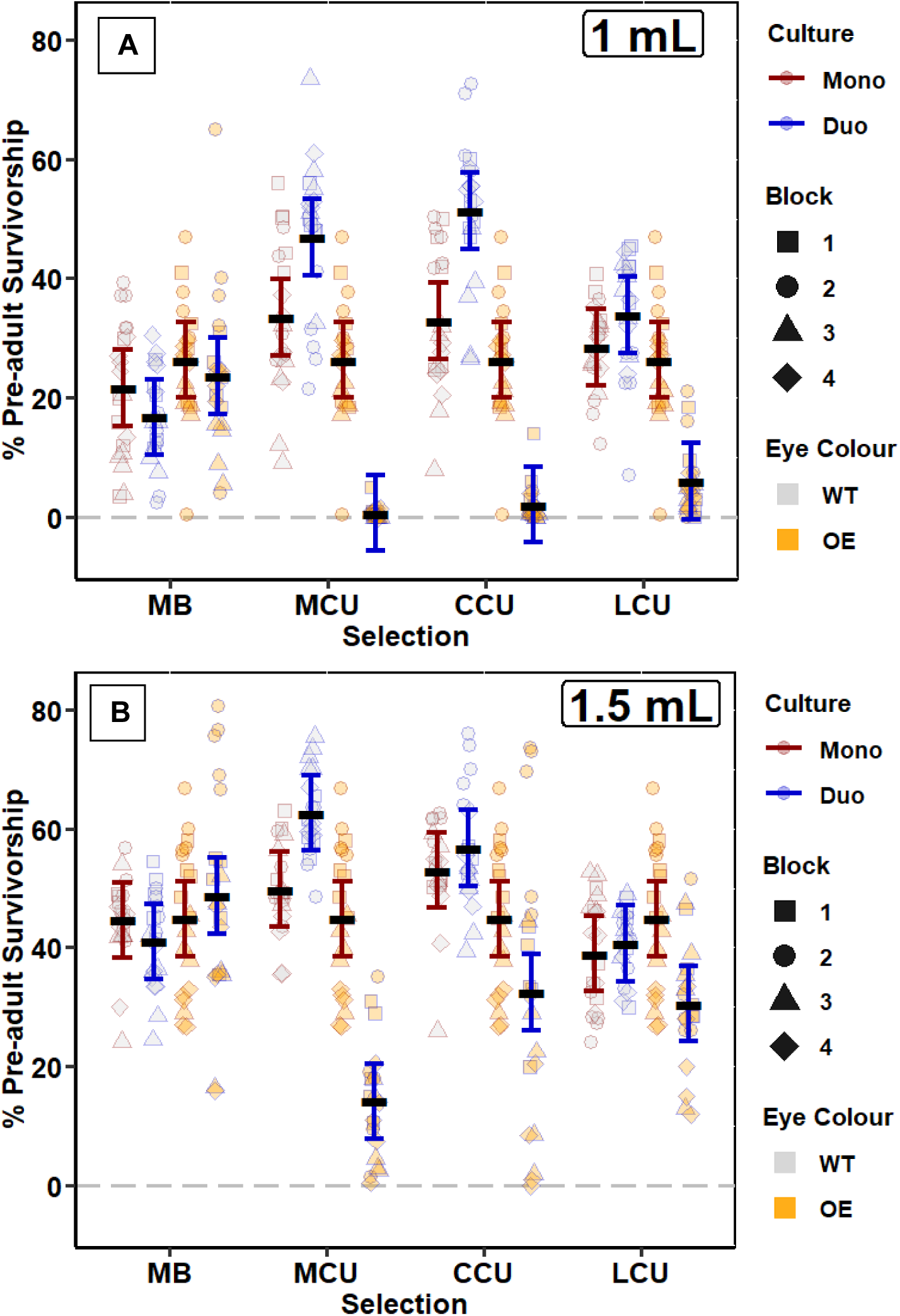
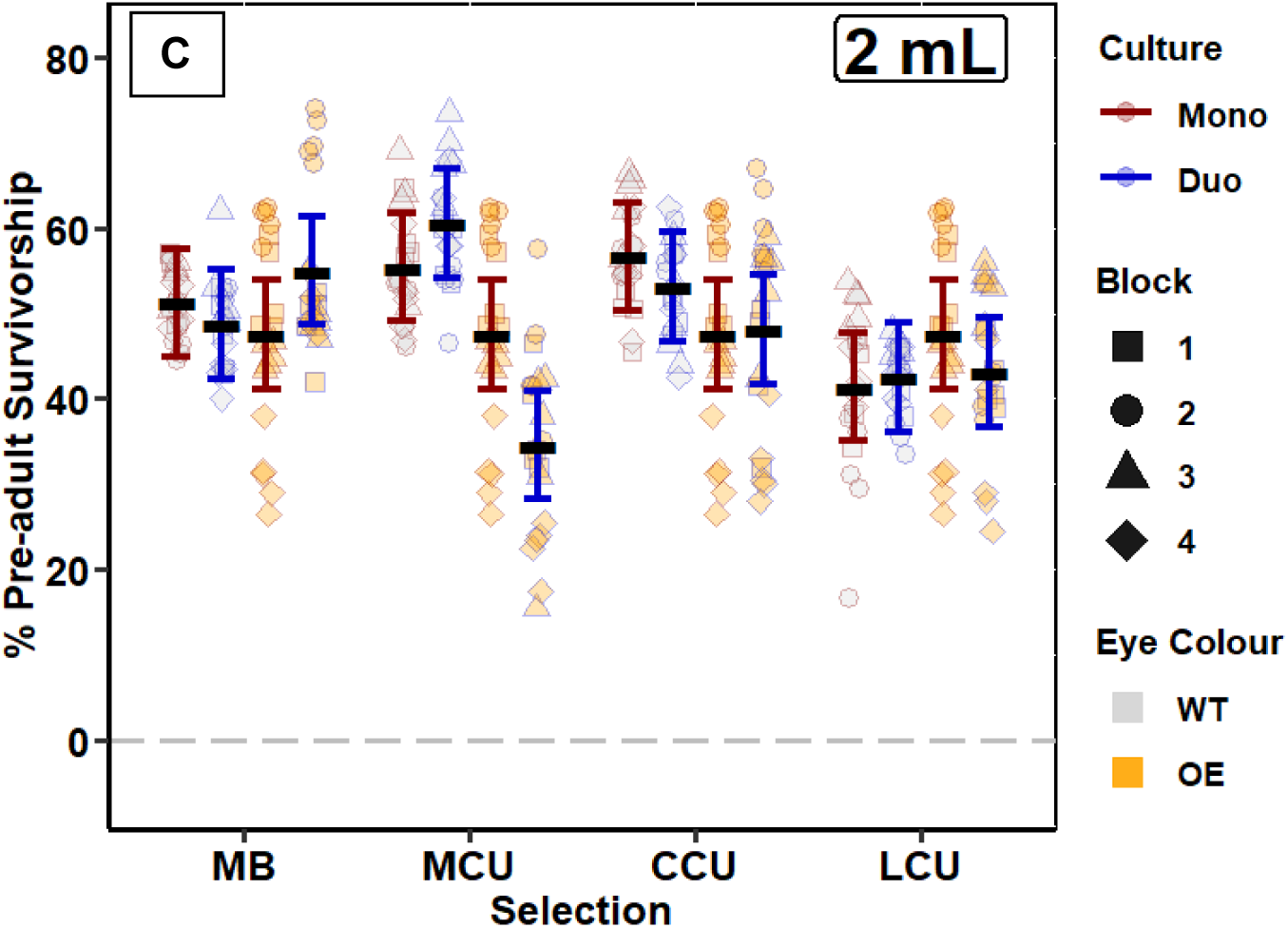
Pre-adult survivorship in A) 1 mL; B) 1.5 mL; C) 2 mL cultures. Black bars in each group represent the mean. Error bars show 95% C.I. for the post hoc test using the relevant within- group error from ANOVA.

**Supp. Fig. 2.**
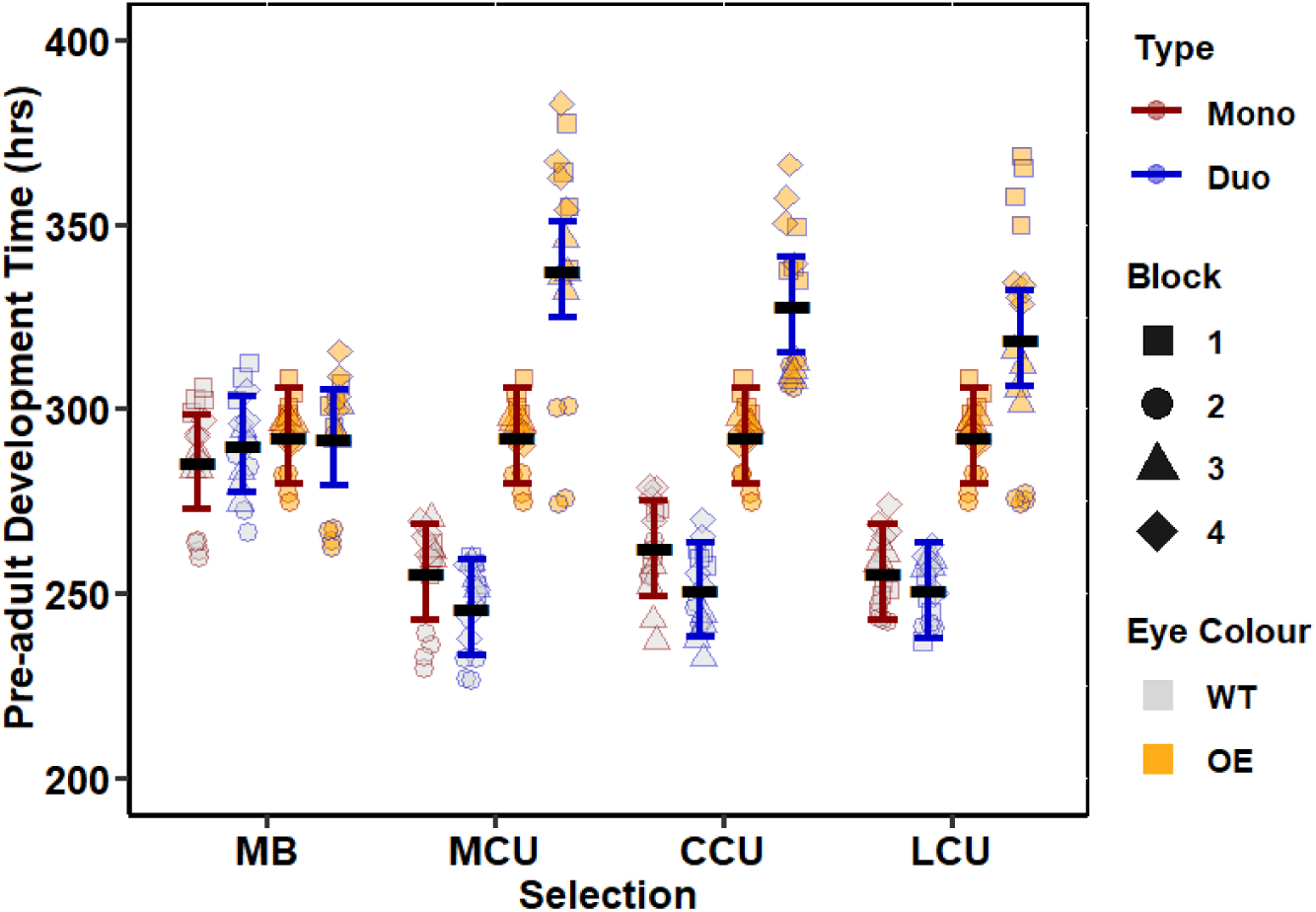
Pre-adult development time (hours) pooling 1.5 mL and 2 mL cultures, as well as male and female data. Black bars in each group represent the mean. Error bars show 95% C.I. for the post hoc test using the relevant within-group error from ANOVA.

**Supp. Fig. 3.**
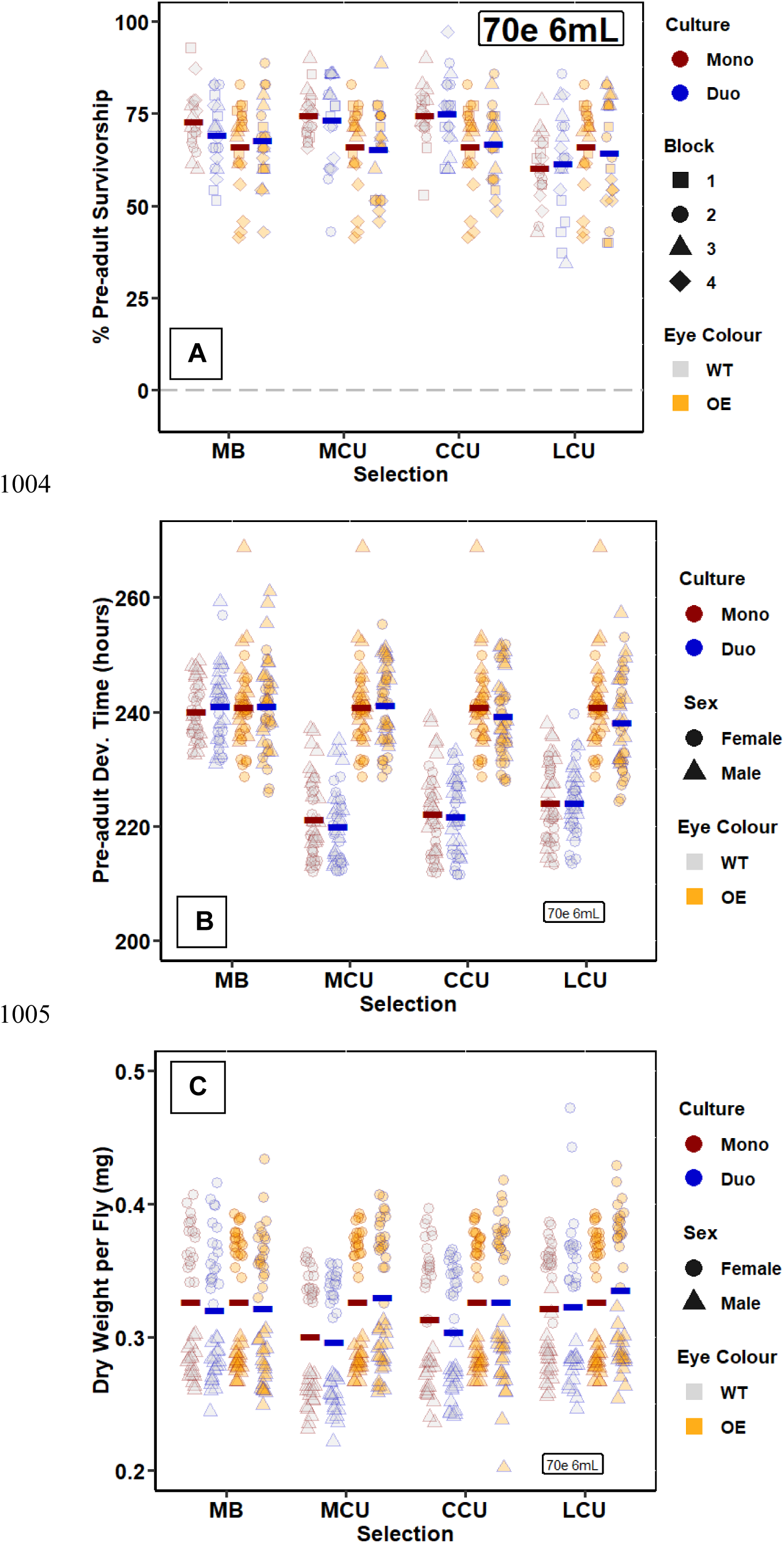
Outcomes at 70 eggs in 6 mL cultures (non-crowded). A) Pre-adult survivorship. B) Pre-adult development time (hrs). C) Dry weight per fly (mg). Coloured bars represent means of each group.

**Supp. Table 1.** ANOVA on tolerance for survivorship.

| Effect | *df* | *df* Error | *F* | *p* |
| --- | --- | --- | --- | --- |
| Selection | 3 | 9 | 2.728070 | 0.106245 |
| Food | 2 | 6 | 1.763133 | 0.249854 |
| Selection*Food | 6 | 18 | 3.149238 | 0.027261 |

**Supp. Table 2.** ANOVA on effectiveness for survivorship.

| Effect | *df* | *df* Error | *F* | *p* |
| --- | --- | --- | --- | --- |
| Selection | 3 | 9 | 24.9151 | 0.000108 |
| Food | 2 | 6 | 38.9512 | 0.000366 |
| Selection*Food | 6 | 18 | 4.1277 | 0.008830 |

**Supp. Table 3.** ANOVA on the raw survivorship data (without effectiveness and tolerance calculations).

| Effect | *df* | *df* Error | *F* | *p* |
| --- | --- | --- | --- | --- |
| Selection | 3 | 9 | 2.1178 | 0.168128 |
| Food | 2 | 6 | 33.7987 | 0.000542 |
| Type | 1 | 3 | 10.6024 | 0.047269 |
| Eye | 1 | 3 | 7.1108 | 0.075910 |
| Selection*Food | 6 | 18 | 4.6312 | 0.005166 |
| Selection*Type | 3 | 9 | 1.8019 | 0.216846 |
| Food*Type | 2 | 6 | 2.2254 | 0.189241 |
| Selection*Eye | 3 | 9 | 43.8494 | 0.000011 |
| Food*Eye | 2 | 6 | 4.2396 | 0.071156 |
| Type*Eye | 1 | 3 | 262.3098 | 0.000512 |
| Selection*Food*Type | 6 | 18 | 0.8149 | 0.572306 |
| Selection*Food*Eye | 6 | 18 | 6.0920 | 0.001263 |
| Selection*Type*Eye | 3 | 9 | 12.2814 | 0.001560 |
| Food*Type*Eye | 2 | 6 | 8.1230 | 0.019620 |
| Selection*Food*Type*Eye | 6 | 18 | 6.1507 | 0.001198 |

**Supp. Table 4.** ANOVA on tolerance for development time.

| Effect | *df* | *df* Error | *F* | *p* |
| --- | --- | --- | --- | --- |
| Selection | 3 | 9 | 7.36425 | 0.008532 |
| Food | 2 | 6 | 3.30806 | 0.107566 |
| Selection*Food | 6 | 18 | 0.84211 | 0.553924 |

**Supp. Table 5.** ANOVA on effectiveness for development time.

| Effect | *df* | *df* Error | *F* | *p* |
| --- | --- | --- | --- | --- |
| Selection | 3 | 9 | 8.052838 | 0.006449 |
| Food | 1 | 3 | 1.407017 | 0.320936 |
| Selection*Food | 3 | 9 | 0.132010 | 0.938569 |

**Supp. Table 6.** ANOVA on the raw development time data (without effectiveness and tolerance calculations).

| Effect | *df* | *df* Error | *F* | *p* |
| --- | --- | --- | --- | --- |
| Selection | 3 | 9 | 2.778 | 0.102499 |
| Food | 1 | 3 | 0.546 | 0.513442 |
| Type | 1 | 3 | 6.427 | 0.085042 |
| Eye | 1 | 3 | 103.760 | 0.002016 |
| Sex | 1 | 3 | 0.002 | 0.969169 |
| Selection*Food | 3 | 9 | 0.531 | 0.672565 |
| Selection*Type | 3 | 9 | 3.750 | 0.053606 |
| Food*Type | 1 | 3 | 3.025 | 0.180370 |
| Selection*Eye | 3 | 9 | 23.034 | 0.000147 |
| Food*Eye | 1 | 3 | 11.999 | 0.040522 |
| Type*Eye | 1 | 3 | 8.600 | 0.060877 |
| Selection*Sex | 3 | 9 | 3.142 | 0.079614 |
| Food*Sex | 1 | 3 | 0.373 | 0.584592 |
| Type*Sex | 1 | 3 | 2.027 | 0.249737 |
| Eye*Sex | 1 | 3 | 4.733 | 0.117847 |
| Selection*Food*Type | 3 | 9 | 0.053 | 0.982687 |
| Selection*Food*Eye | 3 | 9 | 0.105 | 0.954966 |
| Selection*Type*Eye | 3 | 9 | 10.701 | 0.002520 |
| Food*Type*Eye | 1 | 3 | 0.123 | 0.748556 |
| Selection*Food*Sex | 3 | 9 | 0.153 | 0.925020 |
| Selection*Type*Sex | 3 | 9 | 1.842 | 0.209904 |
| Food*Type*Sex | 1 | 3 | 17.001 | 0.025863 |
| Selection*Eye*Sex | 3 | 9 | 2.431 | 0.132202 |
| Food*Eye*Sex | 1 | 3 | 0.879 | 0.417537 |
| Type*Eye*Sex | 1 | 3 | 0.070 | 0.808865 |
| Selection*Food*Type*Eye | 3 | 9 | 0.377 | 0.771984 |
| Selection*Food*Type*Sex | 3 | 9 | 0.103 | 0.956520 |
| Selection*Food*Eye*Sex | 3 | 9 | 0.191 | 0.899642 |
| Selection*Type*Eye*Sex | 3 | 9 | 1.256 | 0.346357 |
| Food*Type*Eye*Sex | 1 | 3 | 2.703 | 0.198687 |
| Selection*Food*Type*Eye*Sex | 3 | 9 | 0.234 | 0.870290 |

**Supp. Table 7.** ANOVA on tolerance for dry biomass.

| Effect | *df* | *df* Error | *F* | *p* |
| --- | --- | --- | --- | --- |
| Selection | 3 | 9 | 6.535786 | 0.012243 |
| Culture | 2 | 6 | 1.851222 | 0.236489 |
| Selection*Culture | 6 | 18 | 4.024106 | 0.009896 |

**Supp. Table 8.** ANOVA on effectiveness for dry biomass.

| Effect | *df* | *df* Error | *F* | *p* |
| --- | --- | --- | --- | --- |
| Selection | 3 | 9 | 20.98280 | 0.000212 |
| Culture | 2 | 6 | 41.31956 | 0.000310 |
| Selection*Culture | 6 | 18 | 1.13016 | 0.384548 |

**Supp. Table 9.** ANOVA on the raw dry biomass data (without effectiveness and tolerance calculations).

| Effect | *df* | *df* Error | *F* | *p* |
| --- | --- | --- | --- | --- |
| Selection | 3 | 9 | 0.7619 | 0.543299 |
| Culture | 2 | 6 | 56.4226 | 0.000129 |
| Culture Type | 1 | 3 | 95.1140 | 0.002290 |
| Time Period | 2 | 6 | 19.5251 | 0.002362 |
| Eye Colour | 1 | 3 | 9.8346 | 0.051823 |
| Selection*Culture | 6 | 18 | 1.8671 | 0.142237 |
| Selection*Culture Type | 3 | 9 | 0.9895 | 0.440505 |
| Culture*Culture Type | 2 | 6 | 0.0926 | 0.912818 |
| Selection*Time Period | 6 | 18 | 6.3127 | 0.001038 |
| Culture*Time Period | 4 | 12 | 20.8850 | 0.000025 |
| Culture Type*Time Period | 2 | 6 | 27.5558 | 0.000946 |
| Selection*Eye Colour | 3 | 9 | 19.9492 | 0.000258 |
| Culture*Eye Colour | 2 | 6 | 0.3575 | 0.713340 |
| Culture Type*Eye Colour | 1 | 3 | 32.9023 | 0.010521 |
| Time Period*Eye Colour | 2 | 6 | 81.5255 | 0.000045 |
| Selection*Culture*Culture Type | 6 | 18 | 1.8061 | 0.154470 |
| Selection*Culture*Time Period | 12 | 36 | 4.9308 | 0.000093 |
| Selection*Culture Type*Time Period | 6 | 18 | 4.2259 | 0.007936 |
| Culture*Culture Type*Time Period | 4 | 12 | 0.6102 | 0.663168 |
| Selection*Culture*Eye Colour | 6 | 18 | 9.7010 | 0.000078 |
| Selection*Culture Type*Eye Colour | 3 | 9 | 10.9354 | 0.002340 |
| Culture*Culture Type*Eye Colour | 2 | 6 | 1.1565 | 0.376003 |
| Selection*Time Period*Eye Colour | 6 | 18 | 18.7577 | 0.000001 |
| Culture*Time Period*Eye Colour | 4 | 12 | 36.6022 | 0.000001 |
| Culture Type*Time Period*Eye Colour | 2 | 6 | 29.7227 | 0.000771 |
| Selection*Culture*Culture Type*Time Period | 12 | 36 | 1.9379 | 0.062373 |
| Selection*Culture*Culture Type*Eye Colour | 6 | 18 | 2.5536 | 0.057390 |
| Selection*Culture*Time Period*Eye Colour | 12 | 36 | 15.4230 | 0.000000 |
| Selection*Culture Type*Time Period*Eye Colour | 6 | 18 | 4.6180 | 0.005237 |
| Culture*Culture Type*Time Period*Eye Colour | 4 | 12 | 8.1356 | 0.002058 |
| Selection*Culture*Culture Type*Time Period*Eye Colour | 12 | 36 | 4.8455 | 0.000109 |

**Supp. Table 10.** ANOVA on tolerance for dry weight per fly.

| Effect | *df* | *df* Error | *F* | *p* |
| --- | --- | --- | --- | --- |
| Selection | 3 | 9 | 1.14488 | 0.382530 |
| Culture | 2 | 6 | 7.10533 | 0.026165 |
| Selection*Culture | 6 | 18 | 1.76215 | 0.163953 |

**Supp. Table 11.** ANOVA on effectiveness for dry weight per fly.

| Effect | df | df Error | F | p |
| --- | --- | --- | --- | --- |
| Selection | 3 | 9 | 8.762882 | 0.004915 |
| Culture | 1 | 3 | 0.463798 | 0.544711 |
| Selection*Culture | 3 | 9 | 2.823193 | 0.099270 |

The following section marks occasions when eclosing flies could not be used for measurement of pre-adult survivorship, pre-adult development time, dry mass at eclosion or dry biomass.

The first section marks ‘general’ exceptions, which were prevalent throughout the culture vials.

The second section marks specific known exceptions, wherein the contents of the particular replicate were discarded due to some issue. In cases where the contents were not discarded, these issues represent some confounding error.

## General Exceptions

f) Some flies were crushed or drowned or escaped. Those were noted down. In case they could be sexed and the eye colour could be identified, they were used for survivorship and development time. None of these flies could be used for weight measurements.
g) Some flies (<5%) in CCU 2 resembled the OE phenotype even under mono-cultures. This was likely to be a preserved mutation in the CCU 2 population and unlikely to be a contamination since CCU flies are maintained under intense selection pressure, under which OE larvae do not survive.
h) Several pupae failed to eclose, or drowned.
i) Freezer switched off for 12 hours from 5-6 March. The temperature was generally cold inside at opening. All flies were tightly sealed in tubes, which were sealed inside two levels of ziplock bags. Thus, there was unlikely to be significant loss of dry mass due to decomposition.
j) Some tube sets were left at room temperature for a few days (tightly sealed in centrifuge tubes, kept inside oven (switched off)). Thus, there was unlikely to be significant loss of dry mass to due to decomposition. This was unavoidable due to COVID-19-related quarantine procedures.

## Specific Exceptions

Block 1:

5. MB 1, 2 mL mono-cultures:
  a. Vials 2 and 5 T2 flies exchanged on 22 to 23 Feb, 2020, reshuffled back with correct numbers. however, the vial identity was lost, thus they could not be used for biomass calculations.
  b. Vials 1 and 4 same as above.
6. LCU 1, 2 mL mono-cultures:
  a. Vials 2 and 3: 3 males, 1 female transferred to vial 2 from vial 3 on 24 Feb, 2020 (T2), this count was reshuffled back. These could not be used for biomass calculations.
7. MCU 1, 1.5 mL mono-cultures: T3, 27 Feb 2020, 3 male and 3 female of 2mL added to T3 tube.

Block 2:

MB 2 1mL duo-cultures 1: vial accidentally frozen on 10 April, 2020. Most eclosions were done. Most pupae had either eclosed or were dried. No further eclosions seen in this vial but also very few eclosions overall in duo-cultures after this.

CCU 2 2mL duo-culture vial 5: frozen on 18^th^ April, 2020. Eclosions were more or less over at this point.

MB 1.5mL duo-culture vial 4: frozen on 14^th^ April, 2020. Eclosions were more or less over by this point.

LCU 2 1.5mL duo-culture: 10^th^ April, 2020 : vials 2, 3 and 4 reshuffled into each other.

Block 3:

1. OE 3 1.5mL, 2mL all vials: 18^th^ march vials added to T2 (should have been T3)
2. MCU 3 2mL duo-cultures: vial 1 and vial 5 reshuffled in T1.

Block 4:

1. LCU 4 1mL mono-culture all vials: first (very few) flies eclosing from all vials were inadvertently discarded.
2. MCU 4 2mL duo-culture: Vial 2 and Vial 5 reshuffled on 6^th^ Feb, 2020. Not used for biomass calculations.

## Notes

### Competing Interest Statement

The authors have declared no competing interest.

